# Metagenome-assembled genomes uncover a global brackish microbiome

**DOI:** 10.1101/018465

**Authors:** Luisa W. Hugerth, John Larsson, Johannes Alneberg, Markus V. Lindh, Catherine Legrand, Jarone Pinhassi, Anders F. Andersson

## Abstract

Microbes are main drivers of biogeochemical cycles in oceans and lakes, yet surprisingly few bacterioplankton genomes have been sequenced, partly due to difficulties in cultivating them. Here we used automatic binning to reconstruct a large number of bacterioplankton genomes from a metagenomic time-series from the Baltic Sea. The genomes represent novel species within freshwater and marine clades, including clades not previously genome-sequenced. Their seasonal dynamics followed phylogenetic patterns, but with fine-grained lineage specific adaptations. Signs of streamlining were evident in most genomes, and estimated genome sizes correlated with abundance variation across filter size fractions. Comparing the genomes with globally distributed aquatic metagenomes suggested the existence of a global brackish metacommunity whose populations diverged from freshwater and marine relatives >100,000 years ago, hence long before the Baltic Sea was formed (8000 years). This markedly contrasts to most Baltic Sea multicellular organisms that are locally adapted populations of fresh- or marine counterparts.

---

Microorganisms in aquatic environments play a crucial role in determining global fluxes of energy and turnover of elements essential to life. To understand these processes through comprehensive analyses of microbial ecology, evolution and metabolism, sequenced reference genomes of representative native prokaryotes are crucial. If these are obtained from isolates, the encoded information can be complemented by phenotypic assays and ecophysiological response experiments to provide insights into the factors that regulate the activity of these populations in particular biogeochemical processes. However, obtaining and characterising new pure cultures is invariably a slow process, even with recent advances in high-throughput dilution culturing approaches ^1^. Most notoriously, the highly-abundant slow-growing oligotrophic lineages typical of pelagic environments ^2,3^ remain severely underrepresented in current culture collections ^4^.

Metagenomics offers a potential shortcut to much of the information obtained from pure culture genome sequencing ^5,6^. The last decade’s revolution in DNA sequencing throughput and cost has provided researchers with the unprecedented possibility of obtaining sequences corresponding to thousands of genomes at a time without the need for isolation, cultivation or enrichment. However, despite vast amounts of sequence data allowing inferences on global distribution of phylogenetic lineages and metabolic potentials ^5–8^, the issue of structuring the data into genomes has remained. This is critical because, while individual genes or genome fragments provide useful information on the metabolic potential of a community, in practice most biochemical transformations take place inside a cell, involving sets of genes structured in controlled pathways. Gaining insight into these pathways requires understanding which genes coexist inside individual microbes. Furthermore, reconstructed genomes from naturally abundant microbes serve as references that allow high-quality annotations to be made in environmental sequencing efforts where otherwise a majority of sequences would remain unclassified ^7,8^. Single-cell sequencing has emerged as a very powerful approach to obtain coherent data from individual lineages ^3,9,10^. This approach allows researchers to select certain targets of interest, based on e.g. cell characteristics or genetic markers, to address specific research questions ^9,11,12^. However, single-cell sequencing requires a highly specialized laboratory facility, and single-amplified genomes (SAGs) typically have low genome coverage, due to the small amount of DNA in each cell and associated whole-genome amplification biases^13^.

While metagenomics can offer, in principle, unlimited amounts of starting material and little amplification bias, it has until recently been impossible to automatically reconstruct full genomes from the mass of genome fragments (contigs or scaffolds) generated from complex natural communities. Approaches based on sequence composition, e.g. tetranucleotide frequencies, have been successfully used to reconstruct near-complete genomes from metagenomic contigs without the use of reference genomes, but can generally only discriminate down to the genus-level ^14,15^. More recently, coverage variation across multiple samples has been used, allowing binning down to species and sometimes strain level ^16–19^. At the same time as genomes are reconstructed, the abundance distribution of these genomes across the samples is obtained, allowing ecological inferences. The CONCOCT (Clustering of contigs based on coverage and composition) program does automatic binning using a combination of these two data sources and was shown to give high accuracy and recall on both model and real human gut microbial communities ^20^, but has not yet been applied to aquatic communities.

The Baltic Sea is, in many aspects, one of the most thoroughly studied aquatic ecosystems ^21^. It presents unique opportunities for obtaining novel understanding of how environmental forcing determines ecosystem structure and function, due to its strong gradients in salinity (North-Southwest), redox (across depths) and organic and nutrient loading (from coasts to center), as well as pronounced seasonal changes in growth conditions. 16S rRNA gene-based studies have revealed prominent shifts in the microbial community composition along these dimensions ^22–24^. The community composition of surface waters changes gradually along the 2000 km salinity gradient, from mainly freshwater lineages in the low salinity North to mostly marine lineages in the higher salinity South-West, and a mixture in the mesohaline central Baltic Sea ^22^. The phylogenetic resolution of 16S amplicons, however, does not permit determining whether prokaryotic lineages are locally adapted freshwater and marine populations or represent distinct brackish strains. A recent Baltic Sea metagenomic study showed how a shift in genetic functional potential along the salinity gradient paralleled this phylogenetic shift in bacterial community composition ^8^. However, since genes were not binned into genomes, different sets of distinguishing gene functions could not be assigned to the genomic context of specific taxa. Reference genomes would therefore be invaluable for a richer exploration of available and future omics data.

Here, we used metagenome time-series data from a sampling station in the central Baltic Sea to generate metagenome-assembled genomes (MAGs) corresponding to several of the most abundant, and mostly uncultured, lineages in this environment. We use these data to compare functional potentials between phylogenetic lineages and relate functionality with seasonal succession. By comparing the MAGs with metagenome data from globally distributed sites, we propose that these are specialised brackish populations that evolved long before the formation of the Baltic Sea and whose closest relatives are today found in other brackish environments across the globe.

## Results and Discussion

### Metagenome-assembled genomes

We conducted shotgun metagenomic sequencing on 37 surface water samples collected from March to December in 2012 at the Linnaeus Microbial Observatory, 10 km east of Öland in the central Baltic Sea. On average, 14.5 million read-pairs were assembled from each sample, yielding a total of 1,443,953,143 bp across 4,094,883 contigs. In order to bin contigs into genomes, the CONCOCT software ^20^ was run on each assembled sample separately, using information on the contigs’ coverages across all samples (Supplementary Fig. 1). Single-copy genes (SCGs) were used to assess completeness and purity of the bins. We approved bins having at least 30 of 36 SCGs present (Supplementary Dataset 1), of which not more than two in multiple copies. This resulted in the identification of 83 genomic bins, hereafter referred to as metagenome assembled genomes (MAGs). The completeness of these MAGs was further validated by assessing the presence and uniqueness of a set of phylum- and class-specific SCGs (Supplementary Dataset 1). Based on these, we estimate the MAGs to be on average 82.7% complete with 1.1% of bases misassembled or wrongly binned. Some MAGs were estimated to be 100% complete. In comparison, recent single-amplified genome studies of free-living aquatic bacteria have obtained average completeness of 55-68% ^3,10^. Importantly, the number of MAGs reconstructed from each sample correlates with the number of reads generated from it and there is no sign of saturation in this trend (Supplementary Fig. 2), meaning that more genomes can easily be reconstructed by deeper sequencing of the same samples. Every sample with over 20 million reads passing quality control yielded at least 3 approved bins. Further, while only highly complete genomes were selected for this study, other research questions might be adequately addressed with partial genomes, many more of which were generated.

In the original CONCOCT study ^20^, binning was done on a coassembly of all samples. Here we employed an alternative strategy where binning was run on each sample separately. This way, community complexity was minimized and binning accuracy increased. Since this strategy may reconstruct the same genome multiple times over the time-series, the 83 complete MAGs were further clustered based on sequence identity. Thirty distinct clusters (BACL [BAltic CLuster] 1-30) with >99% intra-cluster sequence identity were formed (<70% between-cluster identity; 95% sequence identity is a stringent cut-off for bacterial species definition ^25^), that included between one and 14 MAGs each (Supplementary Fig. 3; Table 1; Supplementary Dataset 1). Having several MAGs in the same cluster increases the reliability of the analyses performed, especially in the case of results based on the absence of a sequence, such as missing genes.

**Table 1.** Overview of clusters, sorted by taxonomy.

| Cluster | Num<br>MAGs | Avg bin size<br>(Mb) | Coding<br>(%) | GC<br>(%) | Taxonomy | Abundance<br>avg (max) (%) | Completeness<br>avg (max) (%) |
| --- | --- | --- | --- | --- | --- | --- | --- |
| BACL2 | 7 | 1.07 | 94.3 | 44.3 | Actinobacteria; acl | 1.20 (6.47) | 78.7 (88.2) |
| BACL4 | 6 | 1.01 | 94.8 | 41 | Actinobacteria; acl | 0.36 (1.16) | 80.8 (90.4) |
| BACL15 | 2 | 1.11 | 94.9 | 47.1 | Actinobacteria; acl | 0.37 (1.19) | 84.2 (93.4) |
| BACL6 | 4 | 1.55 | 94.8 | 51.3 | Actinobacteria; aclV | 0.34 (2.92) | 80.0 (86.0) |
| BACL17 | 2 | 1.45 | 95.5 | 52.3 | Actinobacteria; aclV | 0.21 (1.10) | 82.7 (93.4) |
| BACL19 | 1 | 1.26 | 95.3 | 58.1 | Actinobacteria; aclV | 0.09 (0.57) | 59.6 |
| BACL27 | 1 | 1.67 | 94.7 | 50.4 | Actinobacteria; aclV | 0.29 (1.73) | 69.9 |
| BACL25 | 1 | 1.26 | 93.5 | 55.7 | Actinobacteria; Luna | 0.08 (0.62) | 77.2 |
| BACL28 | 1 | 1.12 | 94 | 51.7 | Actinobacteria; Luna | 0.13 (0.96) | 65.4 |
| BACL10 | 3 | 2.57 | 90.2 | 50.5 | Alphaproteobacteria;<br>Rhodobacter | 1.22 (10.91) | 84.0 (88.7) |
| BACL5 | 5 | 1.08 | 96.5 | 30.1 | Alphaproteobacteria;<br>SAR11 | 0.79 (2.92) | 80.2 (89.5) |
| BACL20 | 1 | 1.14 | 95.8 | 30.9 | Alphaproteobacteria;<br>SAR11 | 0.59 (2.56) | 66.9 |
| BACL7 | 3 | 1.74 | 95.4 | 49 | Bacteroidetes;<br>Cryomorphaceae | 0.32 (1.40) | 99.4 (100) |
| BACL11 | 3 | 1.19 | 96 | 32.9 | Bacteroidetes;<br>Cryomorphaceae | 0.43 (1.70) | 75.1 (84.9) |
| BACL18 | 2 | 1.32 | 94.5 | 57.5 | Bacteroidetes;<br>Cryomorphaceae | 0.18 (0.93) | 78.6 (85.7) |
| BACL23 | 1 | 1.73 | 95.3 | 54.6 | Bacteroidetes;<br>Cryomorphaceae | 0.15 (1.45) | 98.3 |
| BACL8 | 3 | 1.74 | 93.6 | 38.7 | Bacteroidetes;<br>Flavobacteriaceae | 0.51 (1.54) | 89.4 (98.3) |
| BACL21 | 1 | 1.92 | 93.1 | 44.1 | Bacteroidetes;<br>Flavobacteriaceae | 0.21 (1.09) | 97.5 |
| BACL22 | 1 | 2.41 | 91 | 32.1 | Bacteroidetes;<br>Flavobacteriaceae | 0.14 (0.98) | 91.6 |
| BACL29 | 1 | 1.48 | 95.2 | 30 | Bacteroidetes;<br>Flavobacteriaceae | 0.08 (0.49) | 88.2 |
| BACL12 | 2 | 2.64 | 93.6 | 47.2 | Bacteroidetes;<br>Sphingobacteriales | 0.20 (2.14) | 85.3 (93.3) |
| BACL14 | 2 | 1.19 | 94.2 | 38.7 | Betaproteobacteria;<br>OM43 | 0.42(1.44) | 86.5 (87.8) |
| BACL30 | 1 | 1.81 | 92.2 | 63.6 | Cyanobacteria;<br>Cyanobium | 0.38 (1.26) | 79.2 |
| BACL3 | 6 | 2.23 | 90.5 | 53 | Gammaproteobacteria<br>; OM182 | 0.80 (3.54) | 85.8 (90.9) |
| BACL1 | 14 | 1.37 | 95.2 | 40 | Gammaproteobacteria<br>; SAR86 | 1.74 (5.23) | 85.9 (90.9) |
| BACL16 | 2 | 2.26 | 91 | 51.3 | Gammaproteobacteria<br>; SAR92 | 0.37 (2.41) | 96.2 (97.2) |
| BACL26 | 1 | 1.91 | 91 | 46 | Gammaproteobacteria<br>; SAR92 | 0.13 (0.70) | 83.2 |
| BACL13 | 2 | 1.01 | 90.9 | 31.9 | Thaumarchaeota;<br>Nitrosopumilaceae | 0.11 (0.88) | 70.0 (78.4) |
| BACL9 | 3 | 1.50 | 93.7 | 56.1 | Verrucomicrobia;<br>LD19 | 0.36 (1.54) | 73.6 (76.4) |
| BACL24 | 1 | 2.98 | 88.4 | 53.1 | Verrucomicrobia;<br>Opitutaceae | 0.08 (2.25) | 92.1 |

The genome clusters generated represent environmentally abundant strains, together corresponding to on average 13% of the shotgun reads in each sample (range: 4 - 23%). This shows that the CONCOCT approach successfully reconstructs novel genomes of environmentally relevant bacteria.

### Phylogeny and functional potential of MAGs

The reconstructed genomes belong to Actinobacteria, Bacteroidetes, Cyanobacteria, Verrucomicrobia, Alpha-, Beta and Gammaproteobacteria and Thaumarchaeota (Table 1, Fig. 1; Supplementary Fig. 4, Supplementary Dataset 2). Phylogenetic reconstruction using concatenated core proteins placed all MAGs consistently within clusters, lending further support to the binning (Supplementary Fig. 4). Based on average nucleotide identity, only BACL8 was estimated to have >70% DNA identity with its nearest neighbour in the phylogenetic tree. In this and many other cases, the closest relative was not an isolate, but a SAG, reflecting these methods’ ability to recover genomes from abundant, but potentially uncultivable, species.

**Figure 1.**
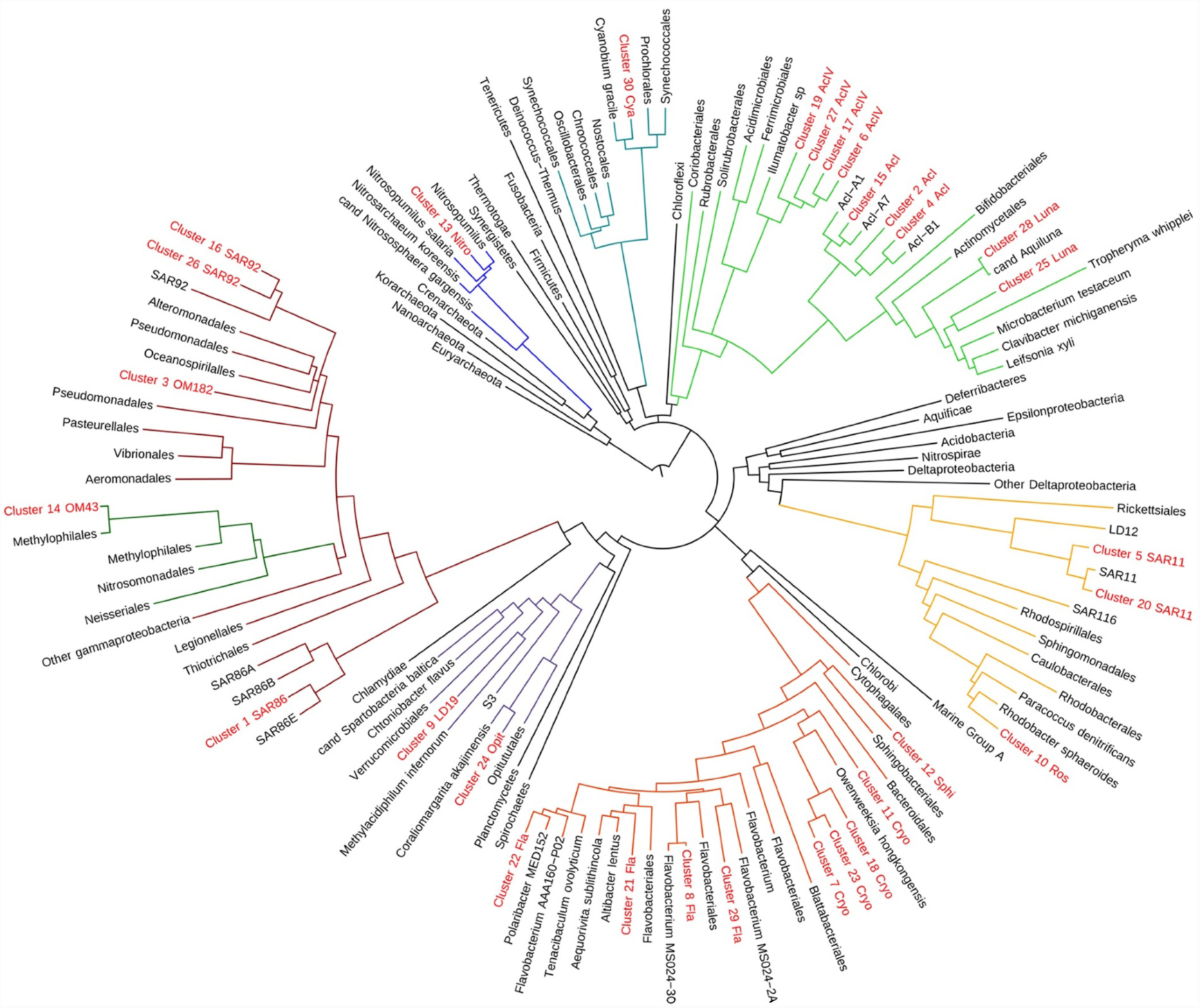
Phylogenetic tree of reconstructed genomes. Phyla and proteobacterial classes for which MAGs were generated are highlighted with coloured branches: Thaumarchaeota (dark blue), Cyanobacteria (blue-green), Actinobacteria (lime green), Alphaproteobacteria (yellow), Bacteroidetes (orange), Verrucomicrobia (purple), Gammaproteobacteria (red) and Betaproteobacteria (dark green).

The broad phylogenetic representation allowed us to compare functional potential between taxonomic groups in this ecosystem. Non-metric multidimensional scaling based on abundance of functional gene categories grouped the MAG clusters according to their phylogeny (Fig. 2; Supplementary Fig. 5; Supplementary Dataset 3), which was confirmed by ANOSIM analysis (Supplementary Table 1). Specifically, Alphaproteobacterial clusters encoded a significantly higher proportion of genes in the “Amino acid transport and metabolism” COG category compared to all other clusters (Welch’s t-test p<0.001). Actinobacteria were significantly enriched in genes in the “Carbohydrate transport and metabolism” COG category (p=0.04). Enzymes involved in carboxylate degradation were significantly more abundant in Gammaproteobacteria compared to all other clusters (p=0.019). Carboxylate degradation enzymes were also abundant in Alphaproteobacteria and Bacteroidetes, but significantly lower in proportion among the Actinobacteria (p<0.01). Bacteroidetes and Verrucomicrobia had the largest number of of glycoside hydrolase genes, including xylanases, endochitinases and glycogen phosphorylases (Supplementary Fig. 6), and thus appear well suited for degradation of polysaccharides such as cellulose, chitin and glycogen, in line with previous findings ^9,26,27^.

**Figure 2.**
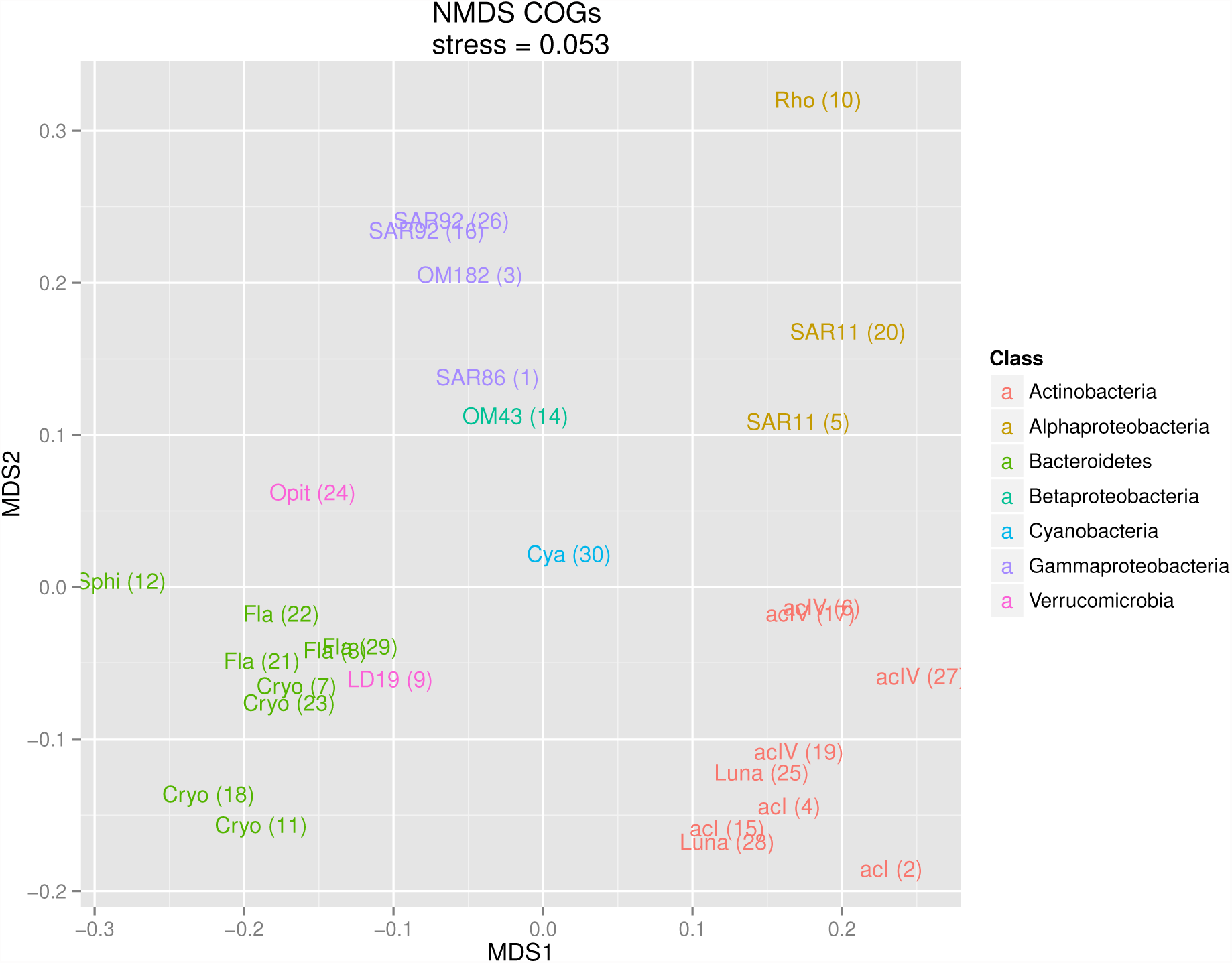
Ordination of MAG clusters based on COG abundances. Non-metric multidimensional scaling was applied to a pairwise distance matrix of the genomes and the first two dimensions are shown. MAG clusters are displayed with abbreviated lineage names and cluster numbers in parentheses and further colored according to Phyla/Class.

Transporter proteins mediate many of the interactions between a cell and its surroundings, thus providing insights into an organism’s niche. A detailed analysis of transporter genes in the 30 MAG clusters (Supplementary Fig. 7; Supplementary Dataset 4) revealed important general patterns, such as a high diversity of genes for amino acid uptake in Actinobacteria and Alphaproteobacteria, a lack of genes for carboxylic acid uptake and a multitude of genes for polyamine uptake in Actinobacteria, and a high diversity of ABC-type sugar transport genes in Actinobacteria and Alphaproteobacteria. The Gammaproteobacteria, Bacteroidetes and Verrucomicrobia encoded a large number of TonB-dependent transporter genes, likely involved in carbohydrate, vitamin and iron chelator uptake ^28^. Phosphate uptake systems, such as the high affinity PstS transporter, were highly abundant in the Betaproteobacterial BACL14, while Thaumarchaeon BACL13 had the highest proportion of phosphonate transporter genes, followed by the Cyanobacterium BACL30 and acIV genome clusters. The BACL30 and the *Nitrosopumilus* BACL13 were sparse in uptake systems for organic molecules in general, consistent with these organisms’ photoautotrophic and chemoautotrophic lifestyles, respectively. In addition to the amoA ammonia monooxygenase gene, BACL13 encoded urease genes, indicating the capability to utilise urea for nitrification, as previously observed in Arctic *Nitrosopumilus* ^29^. In line with the OM43 clade comprising simple obligate methylotrophs with extremely small genomes ^30^, the Betaproteobacterial OM43 cluster (BACL14), which encoded a methanol dehydrogenase gene and genes for formaldehyde assimilation, was also sparse in uptake systems.

All clusters belonging to the Bacteroidetes and Gammaproteobacteria lineages contained the Na+-transporting NADH:ubiquinone oxidoreductase (NQR) enzyme. This enzyme is involved in the oxidative respiration pathway in some bacteria and is similar to the typical H+-transporting ndh NADH dehydrogenase ^31^. However, the NQR enzyme exports sodium from the cell and thereby creates a gradient of Na+ ions, in contrast to the proton gradient generated by the ndh enzyme. The use of the NQR enzyme has been shown to be correlated with salinity (increasing Na+ concentrations) in bacterial communities ^8^. Accordingly, NQR-containing MAG clusters were generally the ones with closest relatives in the marine environment (e.g. Bacteroidetes, see section on biogeography below), while genome clusters more closely related to freshwater bacteria (e.g. Actinobacteria) contained the H+-transporting enzyme. An exception to this were the SAR11 MAGs, which harbored the H+-transporting enzyme despite having predominantly marine relatives. The genomes containing NQR enzymes in our dataset also contained a significantly higher proportion of Na+ symporters and antiporters (for e.g. dicarboxylates, disaccharides and amino-acids), as well as TonB-dependent transporters, compared to the other genomes (Welch’s t-test p<0.001). In contrast, ATP-driven ABC-transporters were significantly less abundant in these clusters (p<0.001), strongly indicating that these bacteria have reduced their energy requirement by making use of the sodium motive force generated by the NQR enzyme to drive transport processes, a strategy that has been suggested previously ^31^. TonB-dependent transporters require energy derived from charge separation across cellular membranes, generally in the form of a proton gradient ^32^. The significant enrichment of TonB transporters in NQR-containing genomes suggests that these proteins may also utilize the sodium motive force.

### Novelly sequenced lineages

The MAG approach has previously proven useful for closing gaps in the tree of life by the reconstruction of genomes from uncultivated species (e.g. ^33,34^). Here we report the first draft genomes for the oligotrophic marine Gammaproteobacteria OM182, and for the typically freshwater Verrucomicrobia subdivision LD19 and Actinobacteria clade acIV. Annotations for these genomes are found in Supplementary Dataset 3 and Supplementary Dataset 4.

OM182 is a globally abundant Gammaproteobacteria which has been grown in enrichment culture, but never sequenced. BACL3 includes a 16S gene 99% identical to that of the OM182 isolate HTCC2188 ^35^. This MAG cluster shares common features with other Gammaproteobacteria, such as a variety of glycoside hydrolases and carboxylate degradation enzymes. It also encodes the cysA sulfate transporter and a complete set of genes for assimilatory sulfate reduction to sulfide and for production of cysteine from sulfide and serine via cysK and cysE. Genes for sulfite production from both thiosulfate (via glpE) and taurine (via tauD) are also encoded in the genome, and this is the only MAG cluster to encode the full set of genes for intracellular sulfur oxidation (dsrCEFH). BACL3 thus appears remarkably well-suited for metabolising different inorganic and organic sulphur sources, the latter potentially originating from phytoplankton blooms ^36^, even more so than previously sequenced isolates of oligotrophic marine Gammaproteobacteria ^37^.

Two verrucomicrobial genome MAG clusters were reconstructed. BACL9 MAGs include 16S rRNA genes 99% identical to that of the globally distributed freshwater clade LD19 ^38^, a subdivision within the Verrucomicrobia still lacking cultured or sequenced representatives. Previous 16S-based analyses placed LD19 as a sister group to a subdivision with acidophilic methanotrophs ^39^. Accordingly, BACL9 is placed as a sister clade to the acidophilic methanotroph *Methylacidiphilum infernorum* ^40^ in the genome tree, but does not encode methane monooxygenase genes and thus lacks the capacity for methane oxidation seen in *M. infernorum*. Interestingly, BACL9 contains a set of genes that together allow for production of 2,3-butanediol from pyruvate (via acetolactate and acetoin). Butanediol plays a role in regulating intracellular pH during fermentative anaerobic growth and biofilm formation ^41^. This is also the only MAG with the genetic capacity to synthesize hopanoid lipids, which have been implicated in enhanced pH tolerance in bacteria by stabilizing cellular membranes ^42^. This indicates adaptation to withstanding lowered intracellular pH such as that induced by fermentative growth under anaerobic conditions. Such conditions occur in biofilms ^41^, and it remains to be shown whether these planktonic bacteria can form biofilms to grow attached to particles in the water column.

BACL 6, 17, 19 and 27 all belong to actinobacterial clade acIV, of the order Acidimicrobiales. Most isolates of the order Acidimicrobiales are acidophilic, and no genomes have been reported for acIV, despite its numeric importance in lake water systems ^43^. Compared to the other typically freshwater clades acI (BACL 2, 4, 15) and Luna (BACL 25, 28), that belong to the order Actinomycetales, acIV MAG clusters have larger genome sizes and contain a significantly lower proportion of genes in the Carbohydrate transport and metabolism COG category (p<0.01), particularly ABC-type sugar transporters (Supplementary Fig. 7). AcIV and acI are also impoverished for phosphotransferase (PTS) genes and amino acid transporters, compared to Luna MAGs. In contrast, acIV MAG clusters contain a significantly higher proportion of genes in the Lipid transport and metabolism COG category (p=0.02), and a significantly higher total proportion of enzymes involved in fatty-acid oxidation (p<0.001), indicating that these Actinobacteria may use lipids as carbon source.

The only cyanobacterial genome assembled was BACL30. While it is placed in the phylogenetic tree as a distant neighbour to *Cyanobium gracile*, its 16S rRNA gene is only 97% identical with it, the same identity as with *Synechococcus* and *Prochlorococcus*. This genome contains genes for the pigments phycocyanin (PC) and phycoerythrin (PE) and harbors the Type IIB pigment gene organization recently identified as being dominant in Baltic Sea picocyanobacteria ^44^. The PC genes cpcBA and the intergenic spacer are 100% identical to sequences in the Type IIB pigment clade. Phylogenies of PC and PE subunits as well as 6 ribosomal proteins consistently placed this cyanobacterial MAG within the Type IIB pigment clades and within the clade of picocyanobacteria whose members are abundant in the Baltic Sea, but for which a reference genome has been unavailable (Supplementary Fig. 8). BACL30 contains the high affinity pstS phosphate transporter, but lacks the phoU regulatory gene as well as an alkaline phosphatase. In this respect the genome is similar to the coastal strain *Synechococcus* CC9311 ^45^, likely reflecting adaptation to higher phosphorous loads compared to the open oceans.

### Genome streamlining and inferred cell sizes

Oligotrophic bacterioplankton are characterised by streamlined genomes, i.e. small genomes with high coding densities and low numbers of paralogs ^46^. For the few cultured oligotrophs, such as *Prochlorococcus* ^47^ and SAR11 ^48^, this coincides with small cell sizes. The small cells render high surface-to-volume ratios, beneficial for organisms that compete for very-low-concentration nutrients ^49^. SAG sequencing has shown that genomic streamlining is a widely distributed feature among abundant bacterioplankton ^3^, contrasting to most cultured marine bacteria. Lauro et al identified genome features for predicting whether an organism or community is oligotrophic or copiotrophic ^2^. Ordination using some of these features (coding density, GC-content and proportion of five COG categories; Swan, 2013) separated our MAG clusters from marine isolate genomes (Fig. 3a). The exceptions were isolates of picocyanobacteria, SAR11 and OM43, that overlapped with our MAG clusters, and the SAR92, OM182 and Opitutaceae MAG clusters that overlapped with the isolates. Hence, most of the MAGs displayed pronounced signs of streamlining. These features, with the exception of GC-content, were found to be highly correlated with genome size (Supplementary Fig. 9), and genome size alone gave equally strong separation (Fig. 3b-c).

**Figure 3.**
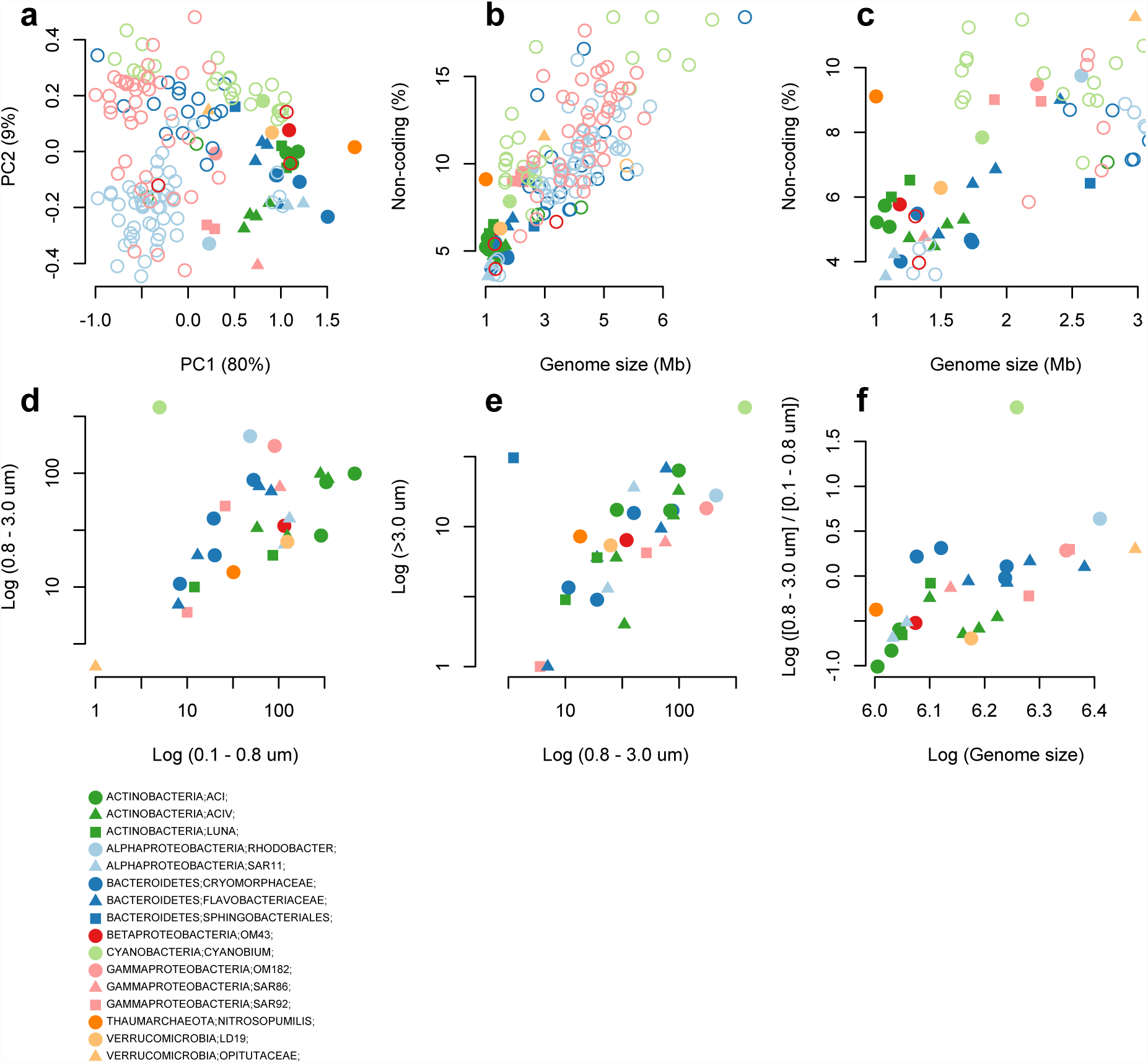
Genome properties and filter size fraction distributions of MAGs. (**a**) Principal Components Analysis (PCA) on our 30 MAG clusters and 135 genomes from marine isolates ^4^ based on log-transformed percentages of non-coding DNA, GC-content, COG category Transcription [K], Signal transduction [T], Defense mechanism [V], Secondary metabolites biosynthesis [Q] and Lipid transport and metabolism [I]. Only isolates belonging to the phyla and proteobacterial classes as represented by MAGs were included. (**b**) Genome size vs. percentage of non-coding DNA plotted for the same set of genomes, (**c**) with a zoom-in on the smaller and denser genomes. (**d-e**) Number of sequence reads matching to our reconstructed genomes from different filter fractions from Dupont et al ^8^. (**f**) The ratio of matches between the 0.8 - 3.0 and the 0.1 - 0.8 um fraction were plotted against genome size, both in log scale.

Interestingly, several of the Bacteroidetes MAG clusters appear to be streamlined, despite Bacteroidetes being generally described as copiotrophic ^46^. One of them (BACL11), which represents a novel branch in the Cryomorphaceae (Fig. 1), has a particularly small genome (1.19 Mbp [range 1.16 - 1.21] MAG size, at 75% estimated completeness) with only 4% non-coding DNA. It encodes a smaller number of transporters than the other Bacteroidetes MAG clusters and only one type of glycoside hydrolase. It also has a comparatively low GC-content (33%). However, the *Polaribacter* MAG cluster (BACL22), which has the largest genome and lowest gene density of the Bacteroidetes genome MAG clusters, has equally low GC-content (32%), as previously observed in planktonic and algae-attached *Polaribacter* isolates ^50^. Since, in general, GC-content correlates very weakly with either genome size or gene density (Supplementary Fig. 9), this may not be an optimal marker for genome streamlining. Supporting the impression that MAGs represent small and streamlined genomes, with little metabolic flexibility, most MAG clusters (25 of 30) encode bacteriorhodopsins (PF01036, Supplementary Dataset 3), which allows them to adopt a photoheterotrophic lifestyle when their required substrates for chemoheterotrophy are not available.

By mapping shotgun reads from different filter fractions (0.1 - 0.8, 0.8 - 3.0 and >3.0 μm) from a previous spatial metagenomic survey of the Baltic Sea ^8^, we could investigate how MAG cluster cells were distributed across size fractions. Comparing counts of mapped reads between the 0.8 - 3.0 and 0.1 - 0.8 fractions showed that Bacteroidetes tended to be captured on the 0.8 μm filter to a higher extent than Actinobacteria (Fig. 3d). This bias could be driven by Bacteroidetes being, to a higher extent, attached to organic matter particles or phytoplankton. However, comparing the >3.0 μm with the 0.8 - 3.0 μm fraction showed a clear bias only for one of the Bacteroidetes clusters (BACL12; Fig. 3e). This cluster has the largest genomes (2.5 and 2.8 Mbp) of the reconstructed Bacteroidetes and is the only representative of the Sphingobacteriales (Fig. 1). Sphingobacteria have previously been suggested to bind to algal surfaces with the assistance of glycosyltransferase genes ^51^. We did not find significantly more glycosyltransferases in BACL12 than in the other Bacteroidetes. Rather, it encodes a greater number of genes containing carbohydrate-binding module (CBM) domains than the other clusters (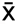 = 12 in BACL12 vs. 1.3 in the other Bacteroidetes and 1.4 in all clusters), which may facilitate adhesion to particles or phytoplankton ^52^.

Since only one Bacteroidetes MAG cluster was biased toward the >3 μm filter, attachment to organic matter doesn’t seem to be the main reason behind the difference in filter capture between Bacteroidetes and Actinobacteria, unless the particles are mainly in the 0.8 - 3.0 um size range. Another possibility is that the bias reflects cell size distributions; each population has a specific size distribution that will influence what proportion of cells will pass through the membrane. Interestingly, the (0.8 - 3.0 um)/(0.1 - 0.8 um) read count ratio is correlated to size of the MAGs (Spearman rho = 0.76; p = 10^−5^; Fig. 3f), indicating a positive correlation between cell size and genome size.

The reason for the streamlining of genomes in oligotrophs is not known ^46^. Lowered energetic costs for replication is one possibility. Despite the energetic requirements for DNA replication being low (<2% of the total energy budget ^53^), the extremely large effective population sizes of oligotrophic pelagic bacteria could explain selection for this trait ^46^. Another possibility is spatial constraints. In *Pelagibacter* the genome occupies 30% of the cell volume ^48^, so that cell size minimisation may be constrained by the genome size. A strong correlation between cell- and genome size for oligotrophic microbes would favor such an explanation. Further analyses with more reconstructed genomes and higher resolution of filter sizes could shed more light on the mechanisms behind genome streamlining.

### Seasonal dynamics

Pronounced seasonal changes in environmental conditions with phytoplankton spring blooms are characteristic of temperate coastal waters. As is typical for the central Baltic Sea, in 2012 an early spring bloom of diatoms was followed by a dinoflagellate bloom, causing inorganic nitrogen to decrease rapidly; later in summer, diazotrophic filamentous cyanobacteria bloomed (Fig. 4 and Supplementary Fig. 10). The only reconstructed picocyanobacteria genome (BACL30) peaked in early summer, between the spring and summer blooms of the larger phytoplankton. A similar pattern was previously observed for an operational taxonomic unit identical to the 16S rRNA gene of the reconstructed genome ^23^.

**Figure 4.**
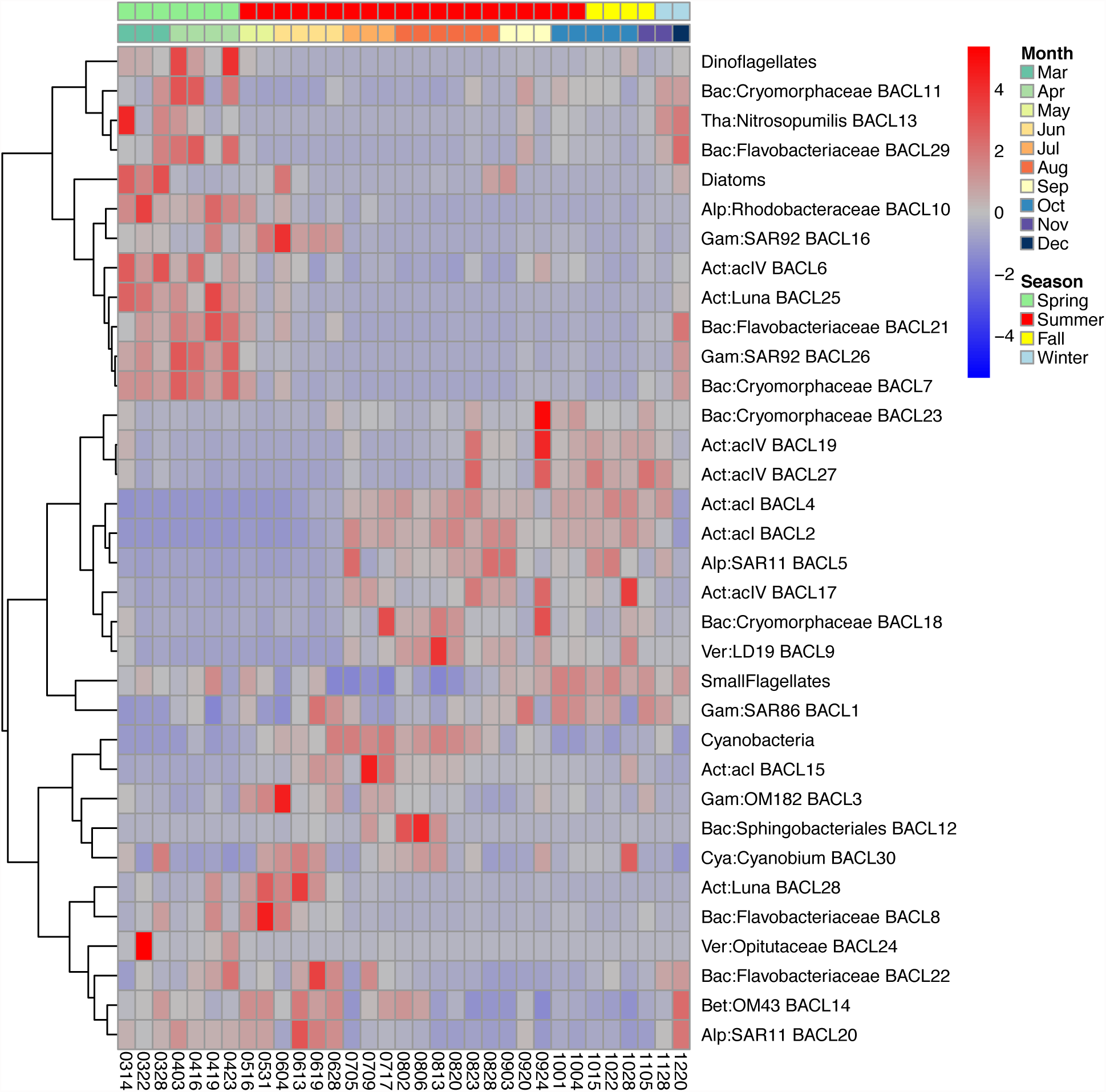
Seasonal dynamics of MAG clusters and phytoplankton. The heatmap plot shows the relative abundance of each MAG cluster in the time-series, based on calculated coverage from read-mappings. In addition, the relative abundance of phytoplankton groups, assessed by microscopy, is shown for the same samples. MAG clusters and eukaryotic groups were hierarchically clustered using Spearman correlations as shown in the dendrogram on the left. Coloured rows at the top indicate month and season of each sample.

The seasonal dynamics of heterotrophic MAGs were highly influenced by the phytoplankton blooms, with different populations co-varying with different phytoplankton (Fig. 4). Phylum-level patterns were present, with a Bacteroidetes-dominated community in spring and early summer (7/9 Bacteroidetes genome MAG clusters), coinciding with the spring phytoplankton blooms, and Actinobacteria being more predominant in the second half of the year (8/10 Actinobacterial genome MAG clusters). This is largely in agreement with what is known for Bacteroidetes: being better adapted to feeding on complex carbohydrates abundant for the duration of phytoplankton blooms ^27^. This was also reflected in the functional annotations, where Bacteroidetes MAGs contained several enzymes for degradation of polysaccharides and were enriched for certain aminopeptidases. For Actinobacteria, no such general correlation pattern to phytoplankton has been shown, but there are indications of association with and active uptake of photosynthates from cyanobacterial blooms ^54,55^. Actinobacteria MAGs, which were enriched in genes for the uptake and metabolism of monosaccharides such as galactose and xylose, became abundant as levels of dissolved organic carbon increased in the water (Supplementary Fig. 10).

Besides these phylum- and order-level trends, temporal patterns were also observed at finer phylogenetic scales. The peaks of Luna clades coincide with phytoplankton blooms, while acI and acIV are more abundant in autumn, after the blooms. AcI also appears to have a more stable abundance profile than acIV, in agreement with Newton et al ^43^. As previously reported for acI SAGs ^10^, cyanophycinase was found in two of the three acI MAG clusters, potentially allowing degradation of the storage compound cyanophycin synthesized by cyanobacteria. The two acI MAG clusters (BACL2 and 4) encoding cyanophycinase became abundant in late July, as filamentous cyanobacteria, which typically produce cyanophycin, started to peak in abundance (Fig. 4). In contrast, all acIV and Luna MAGs lacked this gene.

Furthermore, contrasting dynamics between members of the same clade, as exemplified by one acIV population blooming in spring, highlight that, despite the general similarities in their functional repertoire, lineage specific adaptations allow different microniches to be occupied by different strains (Fig. 2, Fig. 4). As an example, the spring blooming acIV BACL6 contained several genes for nucleotide degradation that were missing in the summer blooming acIV MAG clusters, such as adenine phosphoribosyltransferase, thymidine phosphorylase and pyrimidine utilization protein B. In addition, BACL6 contained genes sulP and phnA for uptake of sulfate and uptake and utilization of alkylphosphonate, respectively. These genes were also found in the spring blooming BACL25 (Luna clade), but were notably absent from the summer blooming acI, acIV and Luna MAG clusters. The capacity to utilize nucleotides and phosphonates as major carbon and phosphorous sources thus probably set BACL6 and 25 apart from other closely related lineages.

The two SAR11 MAG clusters also showed contrasting seasonal patterns, with BACL20 being abundant in spring and peaking in early summer, while BACL5 appeared later and showed a stable profile from July onwards. Functional analysis showed that, of these two populations, BACL5 contained several genes related to phosphate acquisition and storage that were missing from BACL20. These included the high-affinity pstS transporter, polyphosphate kinase and exopolyphosphatase, as well as the phosphate starvation-inducible gene phoH. BACL5 therefore appears better adapted to the low concentrations of phosphate found in mid-to late summer (Supplementary Fig. 10). In addition, proteorhodopsin was found in BACL5, but not in BACL20. However, since the latter consists of only one MAG, this gene may have been missed due to incomplete genome assembly.

### Biogeography of the brackish microbiome

To assess how abundant the MAGs presented here are in other marine and freshwater environments around the globe, fragment recruitment was performed from a collection of samples comprising a wide range of salinity levels. At intermediate levels of sequence identity, different phylogenetic lineages recruit preferentially fresh or marine water fragments. Most markedly, SAR11 displays a clear marine profile, while acI and acIV actinobacteria have a distinct freshwater signature (Fig. 5a, Supplementary Fig. 11a, Supplementary Fig. 12). In addition, MAGs belonging to Bacteroidetes and Gammaproteobacteria show signs of a marine rather than a freshwater signature that fits with the presence of the Na+-transporting NADH dehydrogenase in these lineages. However, at a high identity level (99%) only reads from brackish environments are recruited, including estuaries in North America, to the exclusion of fresh and marine waters much closer geographically to the Baltic Sea (Fig. 5b, Supplementary Fig. 11b, Supplementary Fig. 12). Indeed, it is remarkable that BACL8 is placed phylogenetically as a single clade together with a SAG sampled in the brackish Chesapeake Bay (Supplementary Fig. 4). Despite being separated by thousands of kilometers of salt water, these cells share >92% similarity over Blast+ high-scoring pairs. Overall, our analysis showed that the reconstructed genomes recruited sequences primarily from brackish estuary environments at various levels of sequence identity (Fig. 6).

**Figure 5.**
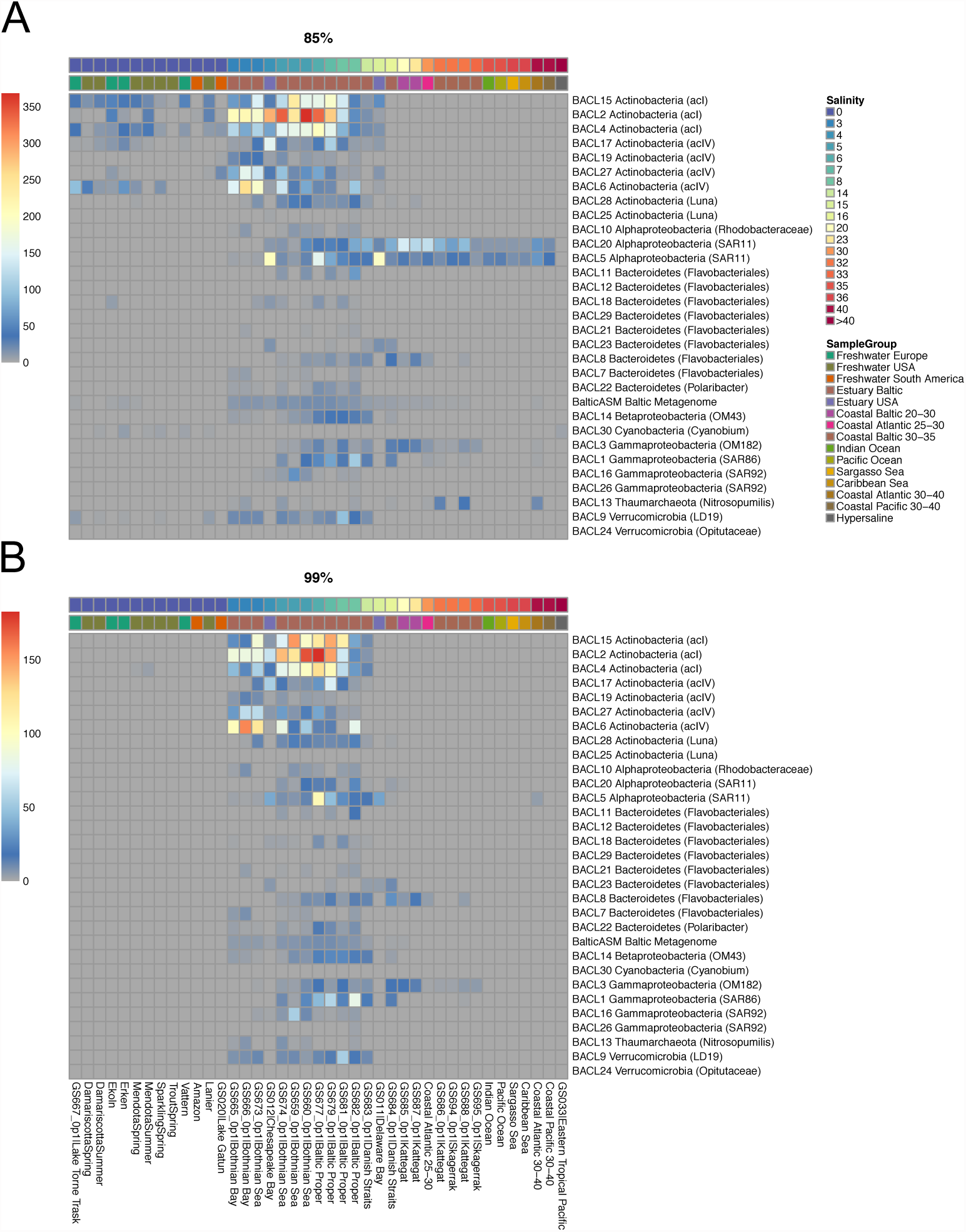
Biogeographical abundance profiles of MAGs. Heatmap plots showing the abundance of recruited reads from various samples and sample groups to each of the 30 MAG clusters at the (**a**) 85% and (**b**) 99% identity cutoff levels. Shown values represent number of recruited reads/kb of genome per 10,000 queried reads. For clarity, several sample groups have been collapsed with recruitment values averaged over samples in the group. Sample groups are indicated by the lower color strip above the plot and samples are ordered by salinity (shown in the upper color strip). See Supplementary Fig. 11 for full visualizations of samples. ‘BalticAsm’ represents a metagenomic co-assembly of all the samples in the time-series.

**Figure 6.**
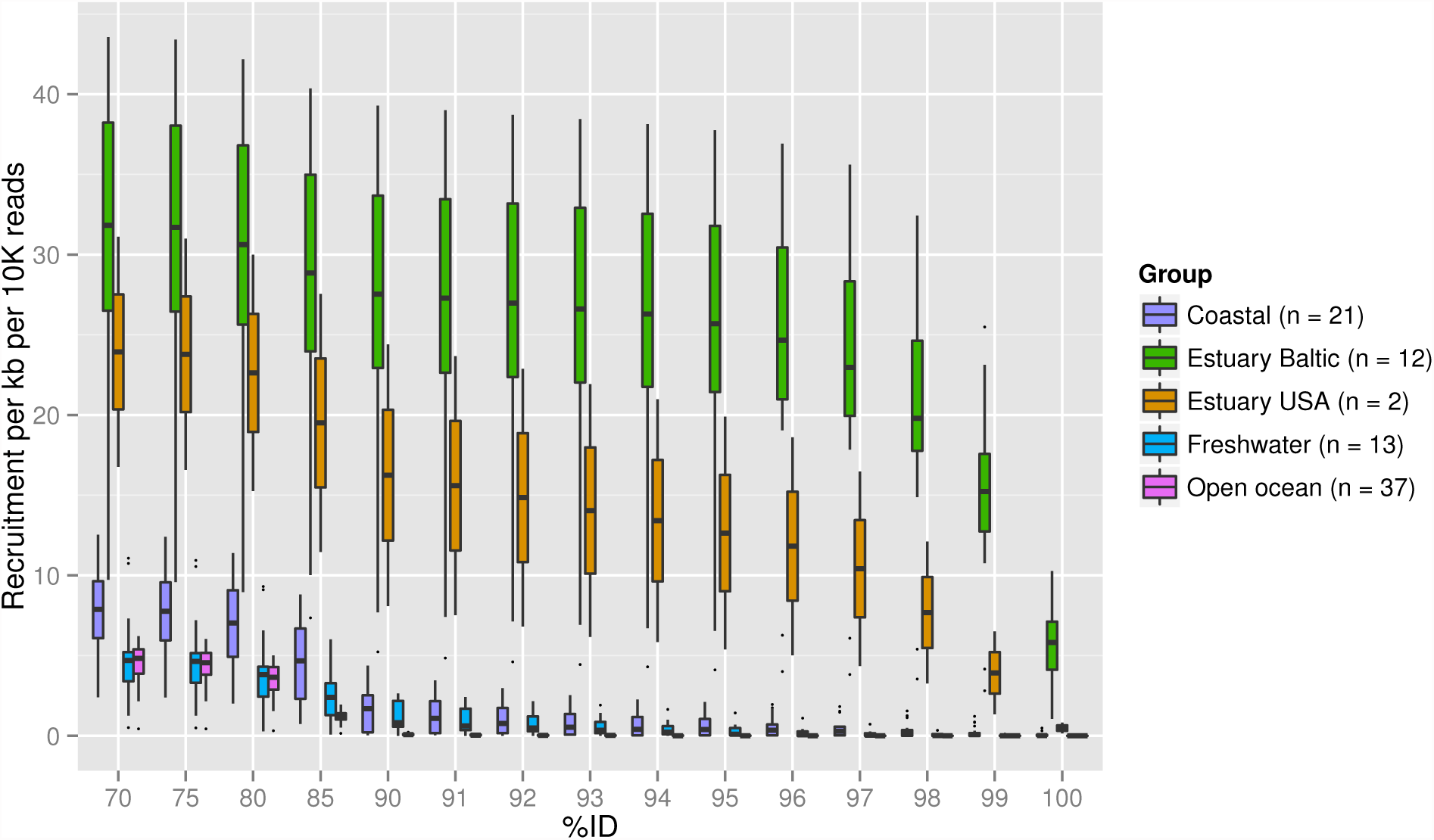
Recruitment profiles at different nucleotide identity. Fragment recruitment values were calculated at various percent identity cutoffs for different sample groups. For each sample in each group, the average recruitment over all 30 MAG clusters was calculated and boxes show the variation of these averages from all samples in each group. The number of samples used in each group are indicated in the legend.

The Baltic Sea is a young system, formed by the opening of the Danish straits to the North Sea in a long process between 13,000 and 8,000 years ago. The initially high salinity has slowly decreased due to the influx of freshwater from the surrounding area and the narrow connection to the open ocean, forming a stable brackish system around 4,000 years ago. Even considering fast rates of evolution for bacteria, the high degree of separation observed at the whole genome level between the Baltic metagenome and global fresh and marine metagenomes cannot be explained by isolation in the Baltic alone. Based on the rates of evolution presented by ^56^, it would take over 100 thousand years for free-living bacteria to accumulate 1% genome divergence. These specialists must therefore have evolved before current stable bodies of brackish water, such as the Baltic, the Black Sea and the Caspian, were formed in the end of the last glacial period. Intriguingly, brackish-typical green sulfur bacteria have been observed in sediment layers of 217,000 years in the now highly saline mediterranean ^57^, suggesting that brackish populations might migrate between these transient environments as salinity shifts. This is in agreement with the well documented separation between freshwater and marine species, which indicates that salinity level is a main barrier isolating populations (reviewed in ^58^). Strains previously adapted to brackish environments and transported through winds, currents or migratory animals can thus proliferate and saturate all available niches before fresh and marine strains can effectively adapt to the new environment.

The prokaryotic populations of the Baltic Sea thus appear to have adapted to its intermediate salinity levels via a different mode than its multicellular species, which are only recently adapted to brackish environments from the surrounding fresh and marine waters ^59,60^. Indeed, while there is low multicellular species-richness and intra-species diversity in the Baltic ^61^, suggestive of a recent evolutionary bottleneck, no such observation has been made for bacteria in the region ^8,22^.

### Conclusion

Here we have presented 83 genomes, corresponding to 30 clusters at >99% nucleotide identity, reconstructed from metagenomic shotgun sequencing assemblies using an unsupervised binning approach. Many of these belong to lineages with no previous reference genome, including lineages known from 16S-amplicon studies to be highly abundant. We show that the seasonal dynamics of these bacterioplankton follow phylogenetic divisions, but with fine-grained lineage specific adaptations. We confirm previous observations on the prevalence of genome streamlining in pelagic bacteria. Finally, we propose that brackish environments exert such strong selection for tolerance to intermediate salinity that lineages adapted to it flourish throughout the globe with little influence of surrounding aquatic communities. The new genomes are now available to the wider research community to explore further questions in microbial ecology and biogeography, solidly placing the automated reconstruction of genomes from metagenomes as an invaluable tool in ocean science.

## Methods

### Sample collection, library preparation and sequencing

Water samples were collected on 37 occasions between March and December of 2012, at 2 m depth, at the Linnaeus Microbial Observatory (N 56° 55.851, E 17° 03.640),10 km off the coast of Öland (Sweden), using a Ruttner sampler. All samples are referred to in the text and figures by their sampling date, in the format yymmdd. Samples were filtered successively at 3.0 μm and 0.22 μm. The 0.22 μm fraction was used for DNA extraction. The procedures for DNA extraction, phytoplankton counts and chlorophyll a and nutrient concentration measurement are described in ^24^.

2-10 ng of DNA from each sample were prepared with the Rubicon ThruPlex kit (Rubicon Genomics, Ann Harbour, Michigan, USA) according to the instructions of the manufacturer. Cleaning steps were performed with MyOne™carboxylic acid-coated superparamagnetic beads (Invitrogen, Carlsbad, CA, USA). Finished libraries were sequenced in SciLifeLab/NGI (Solna, Sweden) on a HiSeq 2500 (Illumina Inc, San Diego, CA, USA). On average, 31.9 million pair-ended reads of 2x100 bp were generated.

### Sequence data quality filtering and assembly

Reads were quality trimmed using sickle (https://github.com/najoshi/sickle) to eliminate stretches where average quality scores fall below 30. Cutadapt ^62^ was used to eliminate adapter sequences from short fragments detected by FastQC (http://www.bioinformatics.babraham.ac.uk/projects/fastqc). Finally, FastUniq ^63^ was used to eliminate reads which were, on both forward and reverse strands, identical prefixes of longer reads (on average, 49% of the reads from each sample). Each sample was then assembled separately, using Ray ^64^ with kmer length 21, 31, 41, 51, 61, 71 and 81. Contigs from each of these assemblies were cut up to 1000 bp in sliding windows every 100 bp using Metassemble (https://github.com/inodb/metassemble) and reassembled using 454 Life Science’s software Newbler (Roche, Basel, Switzerland).

### Binning of sequencing data and construction of MAGs

The quality-filtered reads of each sample were mapped against the contigs of all other samples using Bowtie2 ^65^, Samtools ^66^, Picard (http://broadinstitute.github.io/picard) and BEDTools ^67^. Contigs from each sample were then binned based on their tetranucleotide composition and covariation across all samples using Concoct ^20^ and accepting contigs over 1000, 3000 or 5000 bp in length. Prodigal ^68^ was used to predict proteins on contigs for each bin, and these were compared to the COG database with RPS-BLAST. The resulting hits were compared to a small set of 36 single-copy genes (SCG) used by Concoct, only considering a protein hit if it covered more than half of the reference length. Bins were considered good if they presented at least 30 of the 36 SCG, no more than two of which in multiple copies. Another set of phylum-specific SCG was used to evaluate each selected bin more carefully. Both the general prokaryotic SCG and phylum-specific SCG were selected such that they were present in at least 97% of sequenced representatives within that taxon and had an average gene count of less than 1.03. For the phylum-specific SCG, Proteobacteria was divided down to class level for increased sensitivity.

For each sample, only one CONCOCT run was chosen for downstream analysis. For most samples, the 1000 bp cutoff provided the maximum number of good bins, but samples 120705, 120828 and 121004 had best results with 3000 bp. This resulted in 83 good bins in total. As the same genome could have been independently found in more than one sample, MUMmer ^69^ was used to compare all good bins against each other. The distance between two bins was set as one minus average nucleotide identity, given a minimum of 50% bin coverage of the larger bin in each pair. This procedure yielded 30 clearly distinct clusters (BACL), independently of the clustering method used (average-, full- or single-linkage).

### Abundance estimation and comparison of MAGs

The relative abundance of each MAG was estimated using the fraction of reads in each sample mapping to the respective MAG. Normalized on the size of that bin, this yielded the measure *fraction of reads per nucleotide in bin*. This measure was chosen since it is comparable across samples with varying sequencing output and different bin sizes.

Using the CONCOCT input table, multiplying the average coverage per nucleotide with the length of the contig in question and summing over all contigs within a bin and within a sample gave the number of reads per bin within a sample. The fraction of reads in each sample mapping to each bin was then calculated by dividing this value with the total number of reads from each sample, after having removed duplicated reads.

### Functional analysis

Contigs in each genome cluster were annotated using PROKKA (v. 1.7, ^70^), modified so that partial genes covering edges of contigs were included, to suit metagenomic datasets, and extended with additional annotations so that Pfam (http://pfam.xfam.org/), TIGRFAMs (http://www.jcvi.org/cgi-bin/tigrfams/index.cgi), COG (http://www.ncbi.nlm.nih.gov/COG/) and Enzyme Commission (http://enzyme.expasy.org/) numbers were given for all sequences where applicable. The extended annotation was performed using homology search with RPS-Blast. Metabolic pathways were predicted in MAGs using MinPath (v. 1.2, ^71^) with the Metacyc database (v. 18.1, ^72^) as a reference. Non-metric multidimensional scaling (NMDS) analysis was applied to the genome clusters based on their annotations as well as on a subset of transporter genes and predicted metabolic pathways. Abundances of functional features were explored, and statistical analyses of functional differences between groups of MAGs performed using STAMP (v. 2.0.9, ^73^) with multiple test correction using the Benjamini-Hochberg FDR method.

### Taxonomic and phylogenetic annotation

Initial taxonomy assignment for each MAG was done with Phylosift ^74^. To improve the resolution of annotations, classification of 16S rRNA genes was also used. Complete or partial 16S genes were identified on contigs using WebMGA ^75^. Further, since rRNA is difficult both to assemble and to bin, a complementary approach was used where partial 16S rRNA genes were assembled for each MAG using reads classified as SSU rRNA by SortMeRNA ^76^, but whose paired-end read was assembled in another contig belonging to the same MAG. The identified and reconstructed 16S fragments were classified with stand-alone SINA 1.2.13 ^77^ and by Blasting against the data by Newton et al ^43^.

Using the information provided by Phylosift and 16S analysis, relevant isolate genomes and single-amplified genomes (SAG) were selected. These were combined with all complete prokaryotic genomes in NCBI. Prodigal was used for protein prediction in each genome. These proteomes, together with the proteomes of our MAGs, were used for phylogenetic tree reconstruction using Phylophlan ^78^. Phylophlan’s reference database was not used as we noticed that, in instances where genomes that were already present in the reference were processed by us and added, they tended to branch closer to the MAG than otherwise, thus indicating a role of protein prediction method in the phylogeny. The tree visualizations displayed here were generated with iTOL v2 ^79^. For the sake of clarity, not all species included in the tree are maintained in the overview or clade-specific insets. Since the distance between MAGs and their nearest neighbours in NCBI were as a rule too large for ANI calculation, we adopted Genome BLAST Distance for this comparison ^80^.

### Genome streamlining analysis

The dataset of marine microbial isolates from ^4^ was downloaded from CAMERA (http://camera.calit2.net/). These were functionally annotated in the same way as the MAGs. For streamlining analysis, the GC content, genome length, and average fraction of non-coding nucleotides were calculated. To avoid bias of shorter contigs, the average fraction of non-coding nucleotides was only based on sequences longer than 5000 nucleotides. For clarity, only genomes belonging to the same phyla as our reconstructed MAGs were included in the analysis. For quantifying how MAG cluster cells were distributed across filter size fractions in ^8^, 10,000 random reads were sampled from each size fraction from 21 samples and aligned to the MAGs by BLAST, using 95% identity and alignment length of 100 bp as cutoff.

### Fragment recruitment

Fragment recruitment ^7^ was used to estimate the presence of the reconstructed MAGs in various locations around the globe. We selected a total of 86 metagenomic samples obtained from a wide range of salinity levels and geographic locations (Table 2). Missing salinity value for Delaware Bay (GS011) was set to 15 PSU after consulting the Delaware Bay Operational Forecast System (DBOFS) (http://tidesandcurrents.noaa.gov/ofs/dbofs/dbofs.html).

**Table 2.** Metagenomic projects used as queries for biogeographic fragment recruitments.

| Project | Project ID | Samples | Salinity range (PSU) | Reference |
| --- | --- | --- | --- | --- |
| Global Ocean Sampling Expedition | CAM_PROJ_GOS | 56 | 0.1 - 37, 63 (hypersaline) | 7 |
| Global Ocean Sampling Baltic Sea | CAM_P_0001109 | 19 | 0 - 34 | 8 |
| Freshwater metagenomes | PRJEB4844 | 9 | 0 | 81 |
| Lake Lanier metagenome by 454 | SRR063691 | 1 | 0 | 82 |
| Metagenomics of the Amazon | SRR091234 | 1 | 0 | 83 |

All samples were sub-sampled to 10,000 sequences, each 350 bp in length, and all reads were queried against a database of the reconstructed genome bins using Blast+ (v. 2.2.30). Non-coding intergenic sequences were excluded by using only the nucleotide sequences of predicted open reading frames. Only samples comprising the 0.1-0.8 μm filter fraction were used and only hits with e-value < 1e-5 and alignment length > 200 bp were considered. For visualizations, the number of hits for MAGs in each sample was normalized against the total size (in bp) of the MAG. These normalized counts were then averaged over the MAGs of each MAG cluster.

## Acknowledgements

We thank Anders Månsson and Kristofer Bergström for their knowledgeable and persistent sampling effort, and Sabina Arnautovic and Emmelie Nilsson for their careful processing of samples. This work was funded by the EC BONUS project BLUEPRINT, partially funded by FORMAS, by the Swedish Research Council VR (grant 2011-5689) through a grant to A.F.A, as well as by Formas project ECOCHANGE (Strategic Grant for Marine Research) through a grant to C.L. and J.P. All computations were performed on resources provided by the Swedish National Infrastructure for Computing (SNIC) through the Uppsala Multidisciplinary Center for Advanced Computational Science (UPPMAX) and the PDC Center for High Performance Computing at KTH. Sequencing was conducted at the Swedish National Genomics Infrastructure (NGI) at SciLifeLab in Stockholm.

## Figures and Tables

**Supplementary Dataset 1:** (xlsx-format) For each approved reconstructed bin, the number of bases, number of contigs, taxonomy and copy number of core COGs is reported.

**Supplementary Dataset 2:** (nwk-format) Phylogenetic tree in Newick format including all complete prokaryotic genomes in NCBI ^103^, all approved reconstructed bins and selected single amplified genomes and isolate genomes.

**Supplementary Dataset 3:** (xlsx-format) Presence or absence of every COG/PFAM/TIGRFAM and enzyme in each approved bin.

**Supplementary Dataset 4:** (xlsx-format) Transporter genes included in each gene cluster.

**Supplementary Figure 1.**
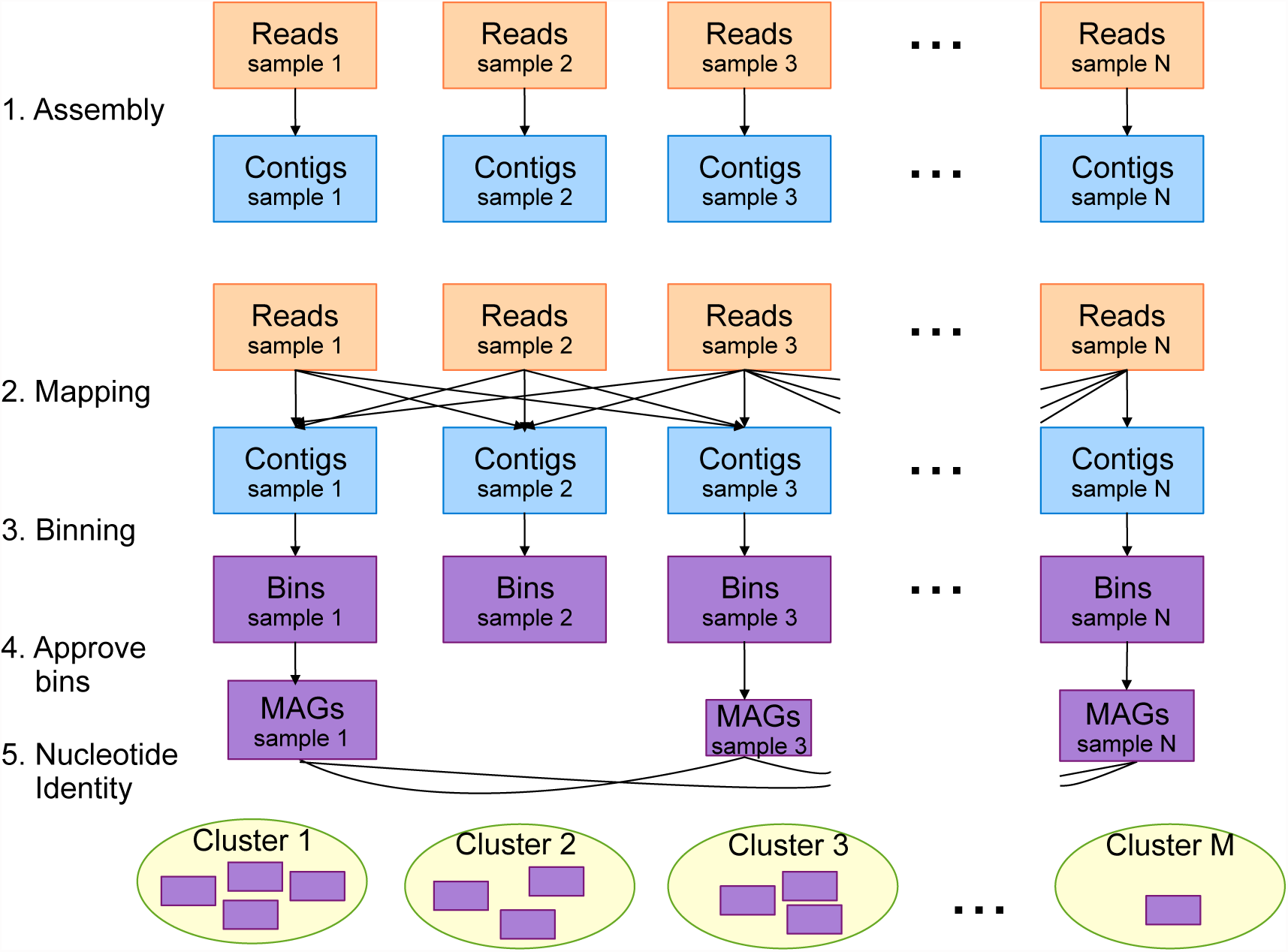
Overview of the genome reconstruction procedure. 1) Reads from each sample are assembled separately. 2) Reads from each of the samples are mapped onto the contigs of all samples, generating a coverage profile for each contig. 3) Together with tetranucleotide composition, these profiles are used to bin contigs in each sample with CONCOCT. 4) Completeness and purity of the bins are evaluated using single-copy housekeeping genes, and approved bins are selected. 5) Approved bins (Metagenome-assembled genomes; MAGs) from all samples are cross-compared using sequence identity and highly similar (>99% identity) MAGs are clustered.

**Supplementary Figure 2.**
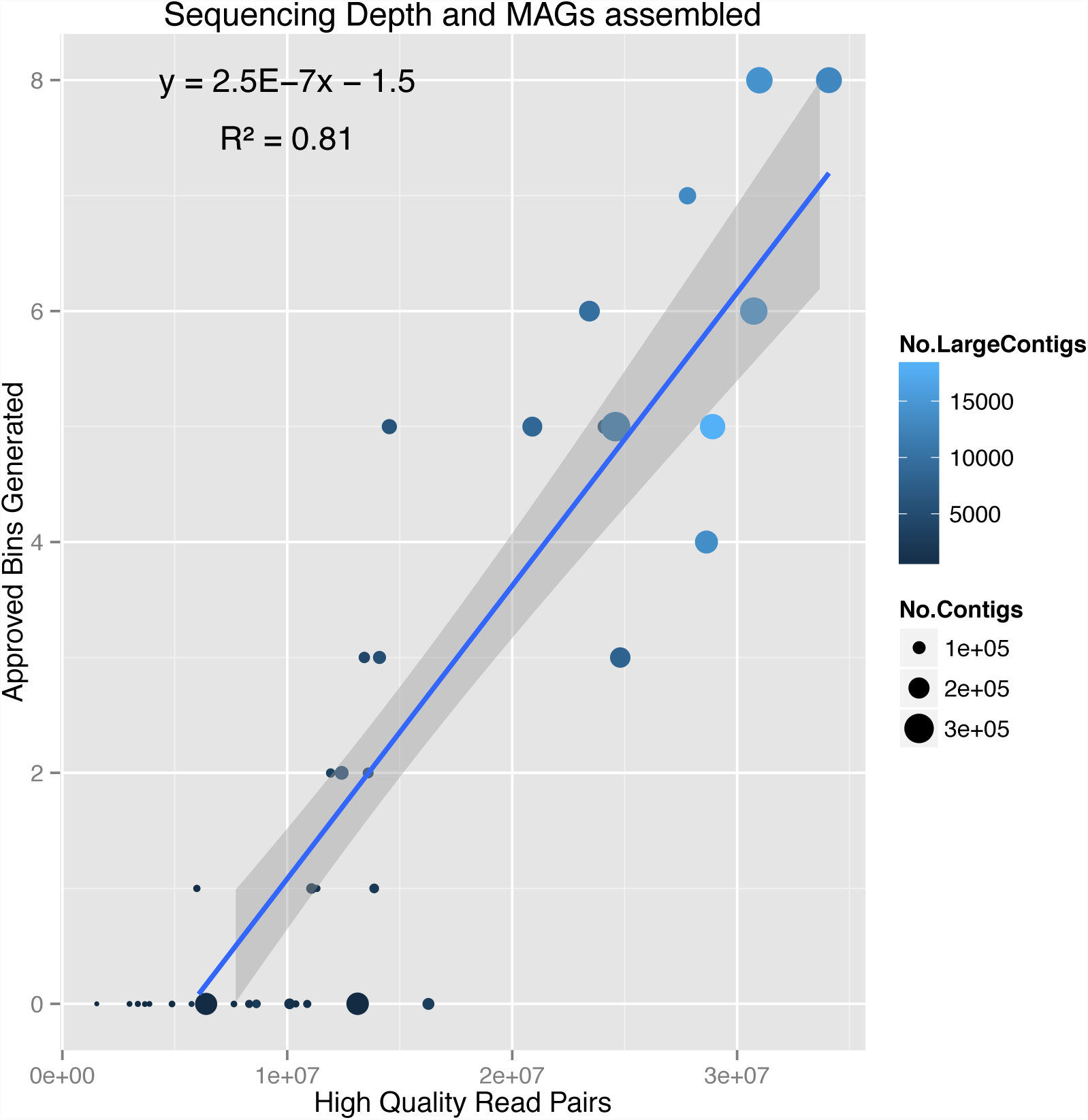
The number of approved bins generated from each sample is directly proportional to the number of sequencing reads passing quality control, the number of contigs assembled and the number of contigs of at least 1000 bp (LargeContigs).

**Supplementary Figure 3.**
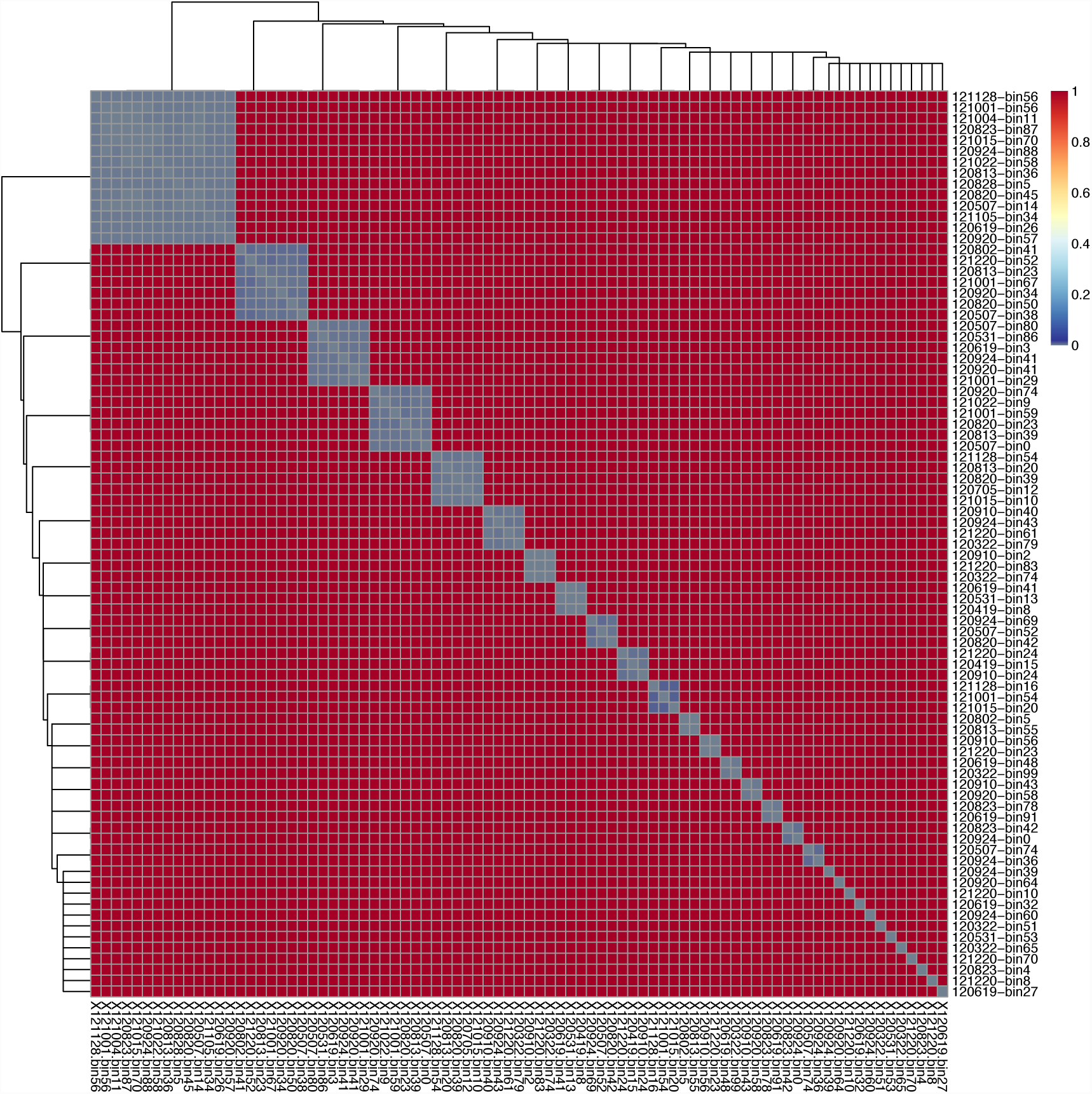
Average-linkage clustering of MAGs. Distance is defined as 1 minus the average nucleotide identity between MAGs, or set to 1 when the coverage of the larger MAG over the smaller is less than 50%.

**Supplementary Figure 4.**
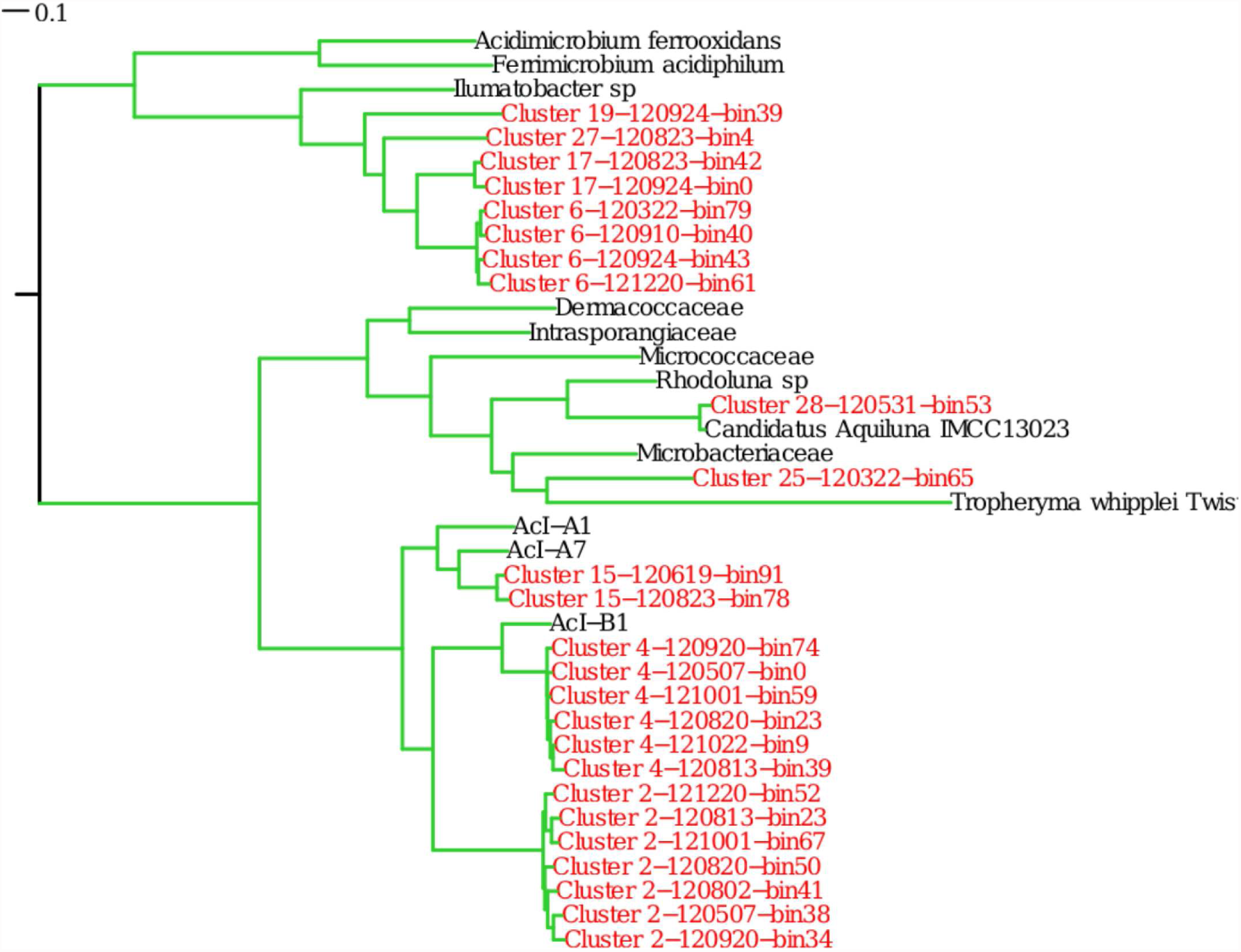
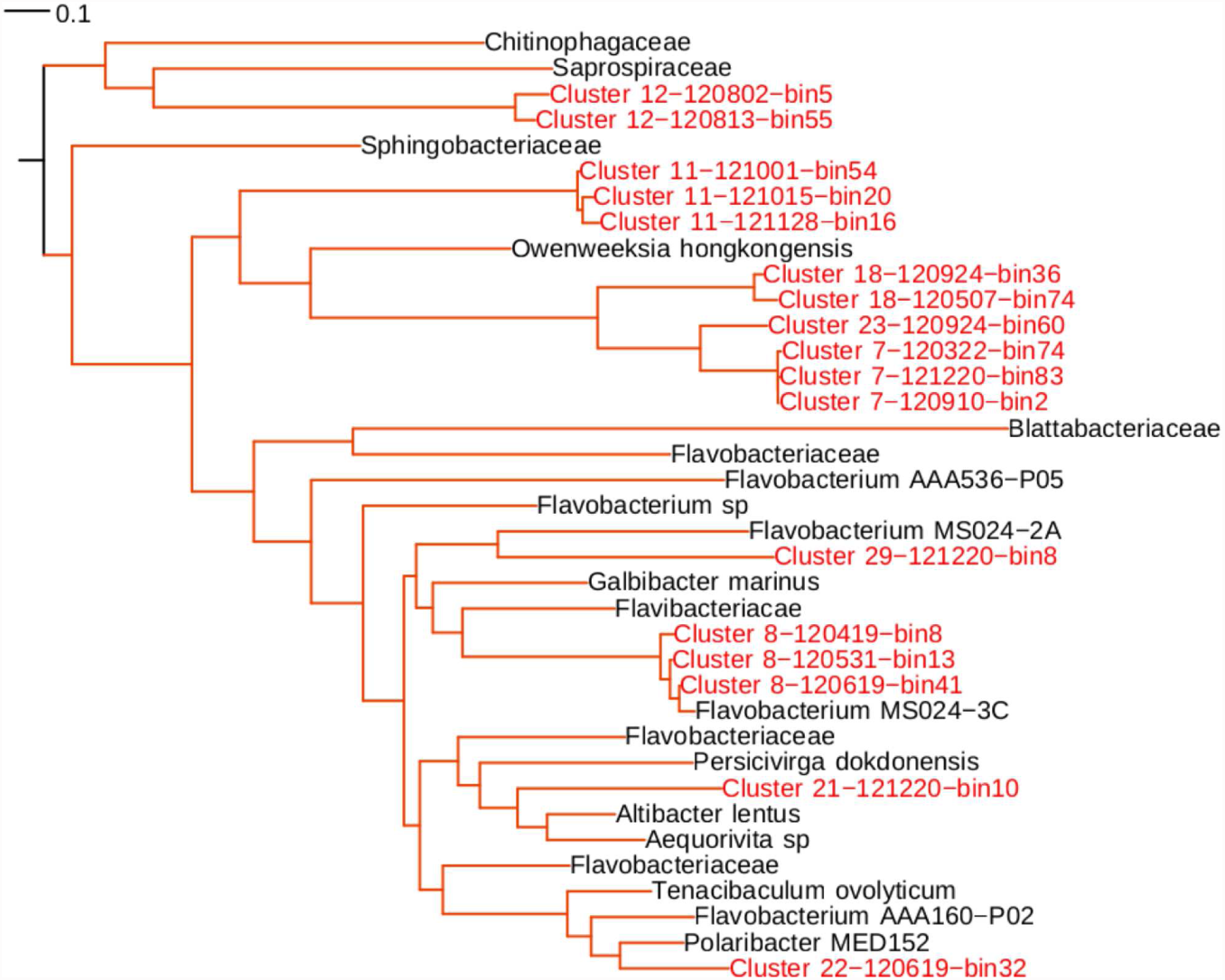
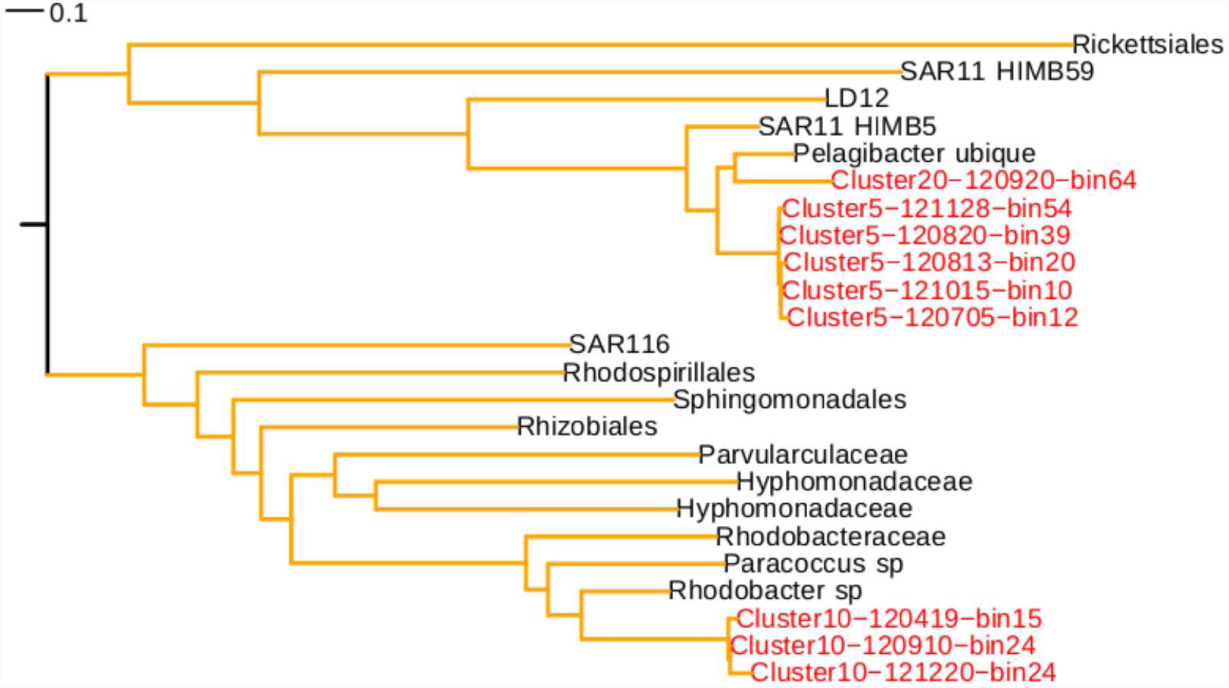
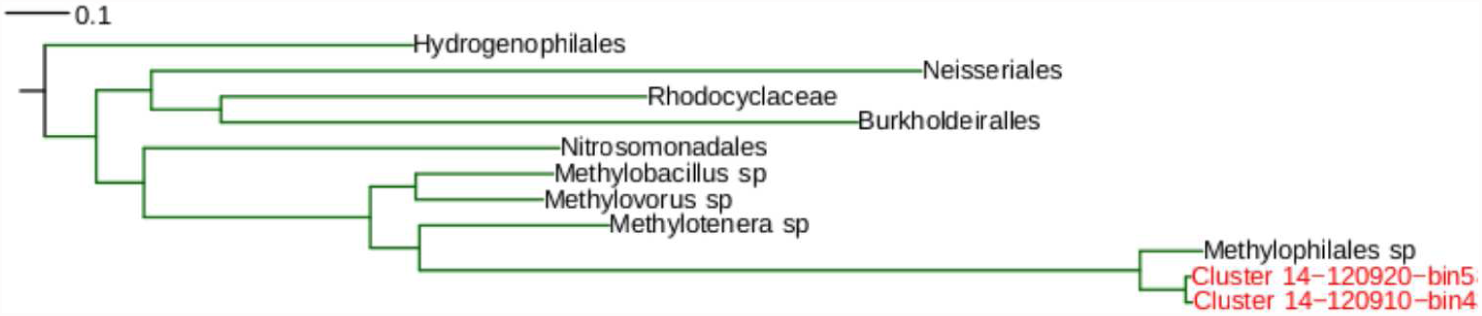
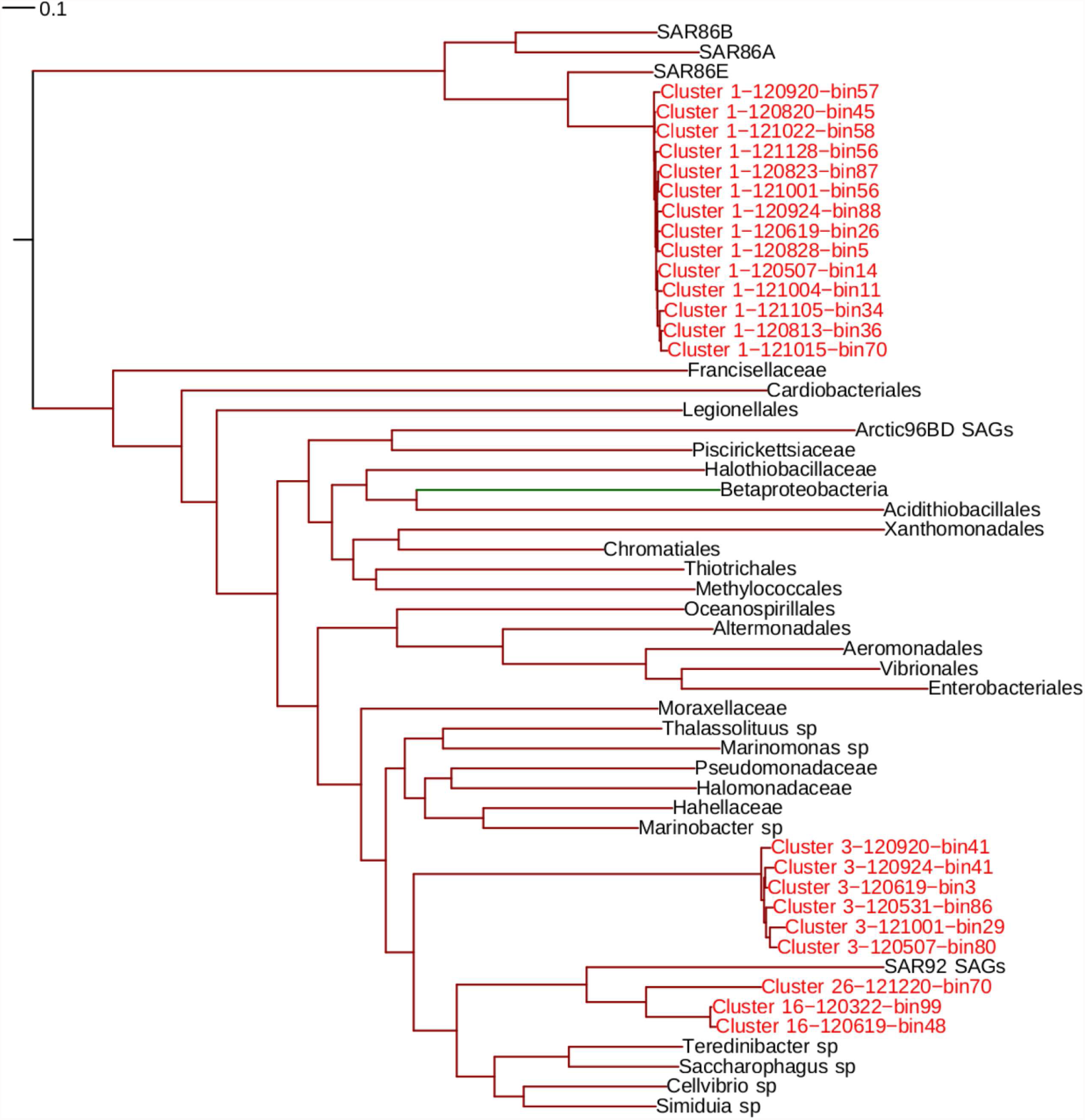
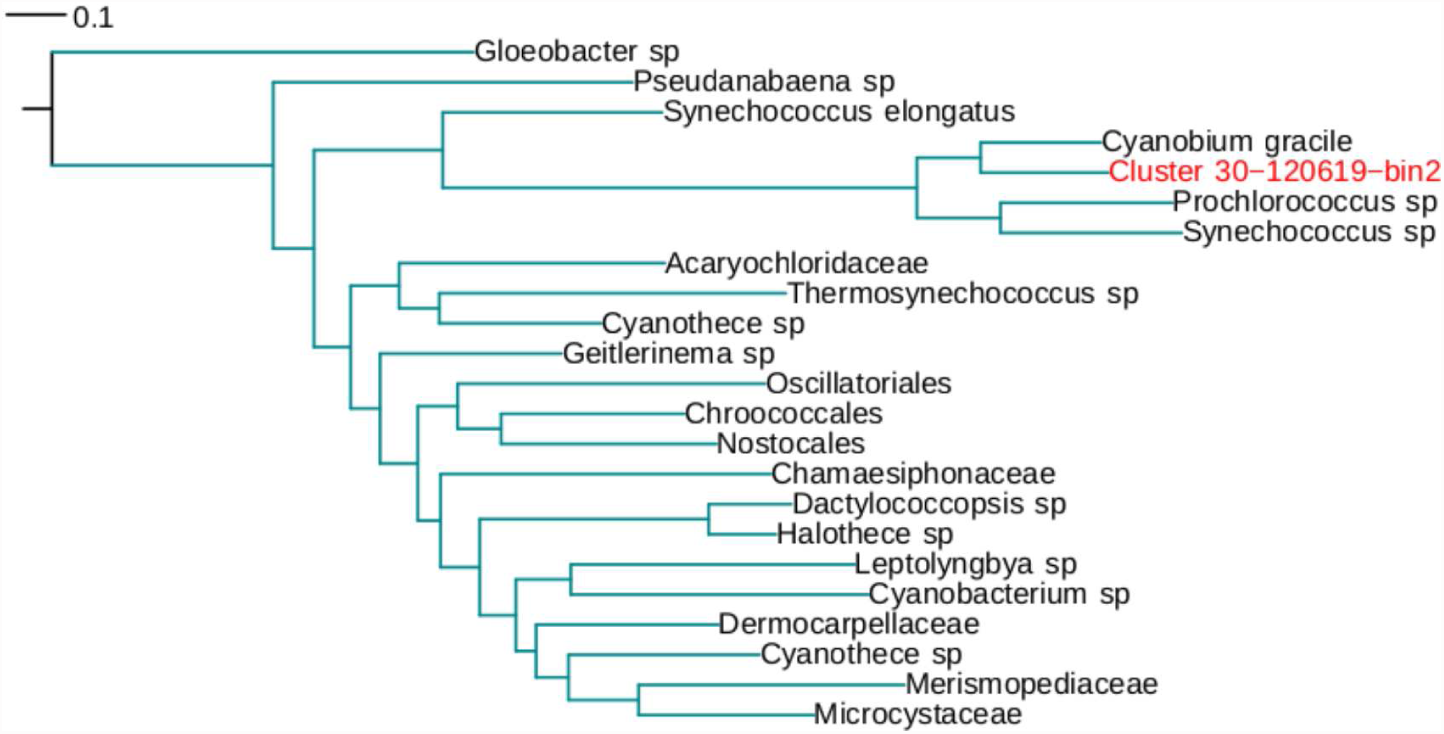
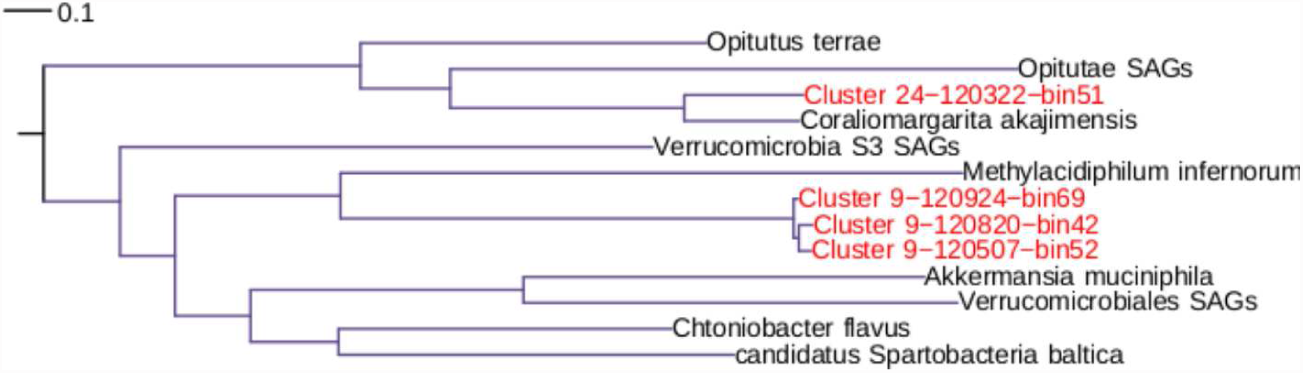
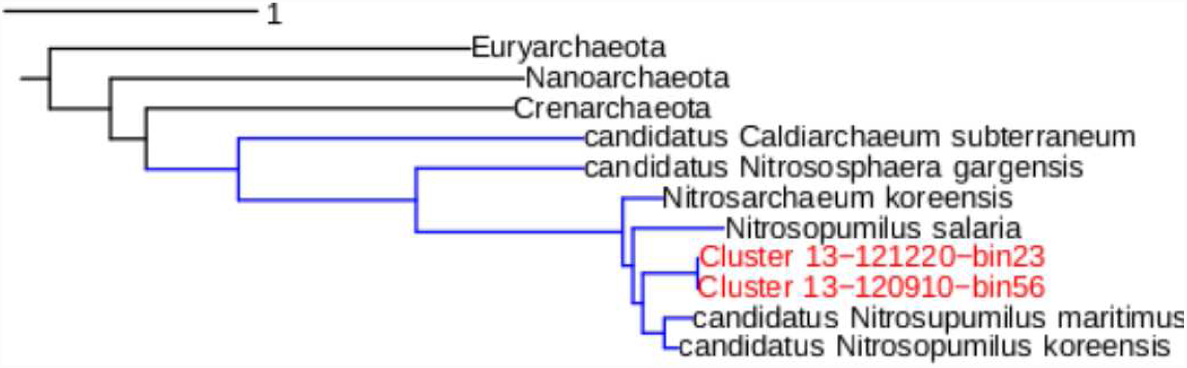
Phylogenetic placement of the MAGs generated in different phyla/classes. (**a**) Actinobacteria. Reconstructed actinobacterial genomes belong to the lineages acI, acIV and Luna. Ac-I and Luna belong to the order Actinomycetales, while ac-IV belong to Acidimicrobiales. Isolates of the Luna clade have had their genome sequenced ^84,85^. Isolation of acI and acIV has been unsuccessful, but SAGs have recently been published for acI-A and acI-B ^10,86^. Reconstructed 16S fragments placed MAGs in lineages acI-A, acI-B, acI-C, acIV-A, acIV-C and Luna ^43,87–90^. Consistent with previous estimates based on a few contigs identified as acIV 83 these MAGs are relatively low GC (average 53%). (**b**) Bacteroidetes. The reconstructed Bacteroidetes genomes belong to the families Flavobacteriaceae, Cryomorphaceae and the order Sphingobacteriales. Cryomorphaceae is a poorly studied family with just three isolate genomes and one SAG available ^91^, not closely related to the clusters that form three major branches presented here. (**c**) Alphaproteobacteria. SAR11 and Rhodobacteraceae clusters were constructed. The two Pelagibacteriaceae clusters are clearly placed in the marine Ia clade, contrary to a previous study that suggested Baltic Pelagibacteriaceae belonged to the brackish clade IIIa or the freshwater clade LD12 ^8^. (**d**) Betaproteobacteria. One OM43 MAG was constructed. Up to now, only two reference genomes were available for this methylotrophic clade, both from the Pacific Ocean ^30,92^ (**e**) Gammaproteobacteria. Known marine clades of Gammaproteobacteria were reconstructed, including SAR86, SAR92, and OM182. No reference genome was previously available for the latter. (**f**) Cyanobacteria. The cyanobacteria genome reconstructed here belongs to a picocyanobacteria with 100% 16S identity to various freshwater *Synechococcus/Cyanobium* ^93^. An operational taxonomic unit of identical 16S has also been observed to be abundant across the Baltic with a strong spatial correlation with the previously described Verrucomicrobia MAG “Candidatus Spartobacteria baltica” ^22^. (**g**) Verrucomicrobia. The Verrucomicrobia phylum has been divided into five monophyletic subdivisions ^94^, but shortly afterwards a novel freshwater subdivision (LD19; ^95^) and a subdivision consisting of acidophilic methanotrophs ^96^ were found. BACL24 is phylogenetically placed within the family Opitutaceae (Subdivision 4), closest to the freshwater isolate *Coraliomargarita akajimensis ^97^*. BACL9 forms a sister branch to *Methylacidiphilum infernorum*, but their 16S rRNA genes are only 84% identical. Instead, 16S positions BACL9 as a member of LD19. (**h**) Thaumarchaeota. Thaumarchaeota are very abundant in the global ocean ^98^, where they play important roles in the nitrogen and carbon cycles by driving ammonia oxidation ^99^. A single archaeal genome cluster was reconstructed, whose 16S rRNA gene is 99% identical to marine lineages of *Nitrosopumilus maritimus ^100,101^*). *N. maritimus* are known to be abundant in Baltic Sea suboxic waters and this MAG has 98% identity to the previously described Baltic *N. maritimus* GD2 (98%; ^102^). However, a whole genome comparison does not support the placement of these MAGs in the same species as *N. maritimus*, but rather in the same genus or family.

**Supplementary Figure 5.**
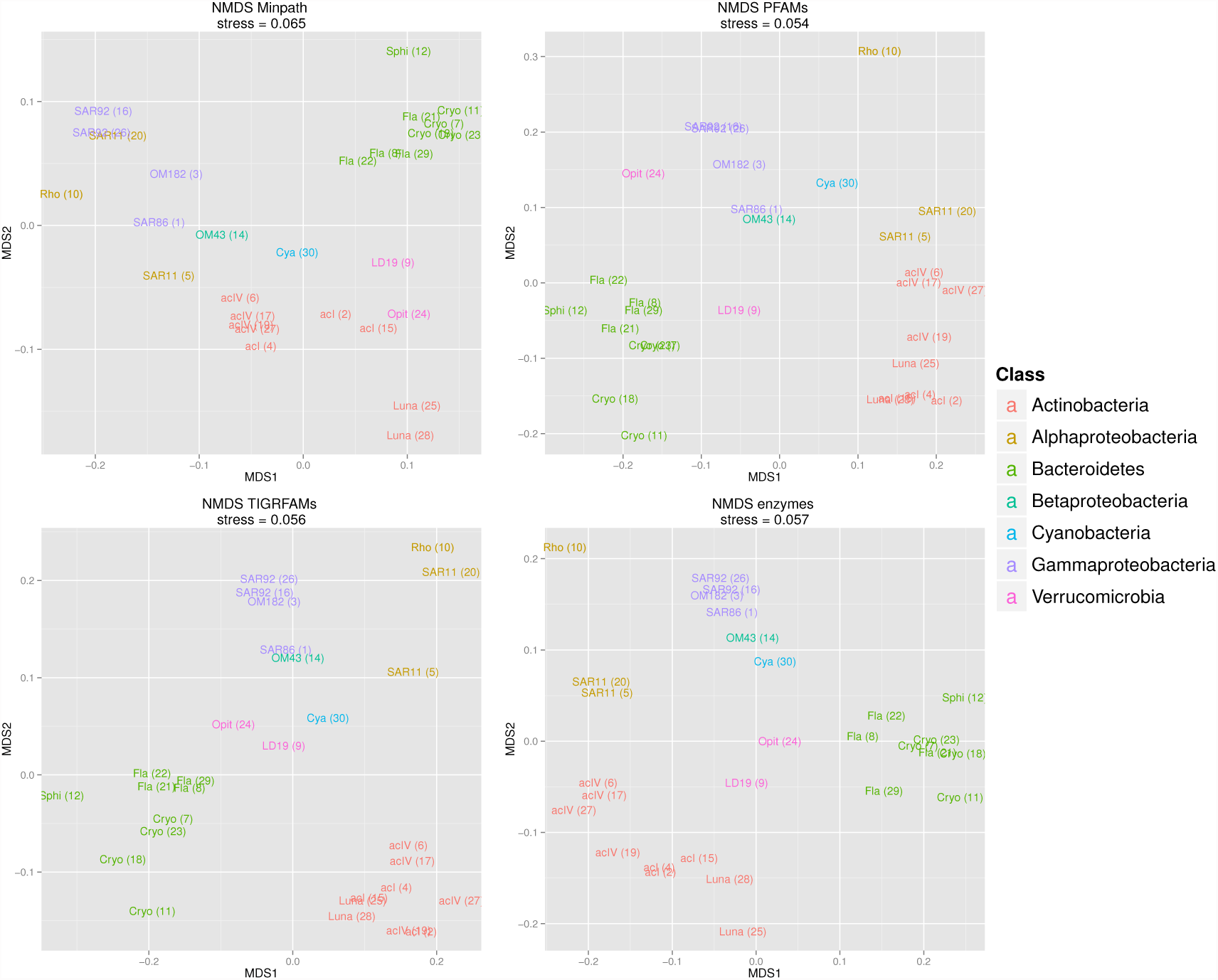
Non-metric multidimensional scaling plots based on functional annotations. Non-metric multidimensional scaling was applied to a pairwise distance matrix of the genomes and the first two dimensions are shown. Genome clusters are displayed with abbreviated lineage names and cluster numbers in parentheses and further colored according to Phyla/Class. Based on enzyme and metabolic pathway predictions (enzyme and Minpath, respectively) and hidden Markov model profile designations (PFAM and TIGRFAM).

**Supplementary Figure 6.**
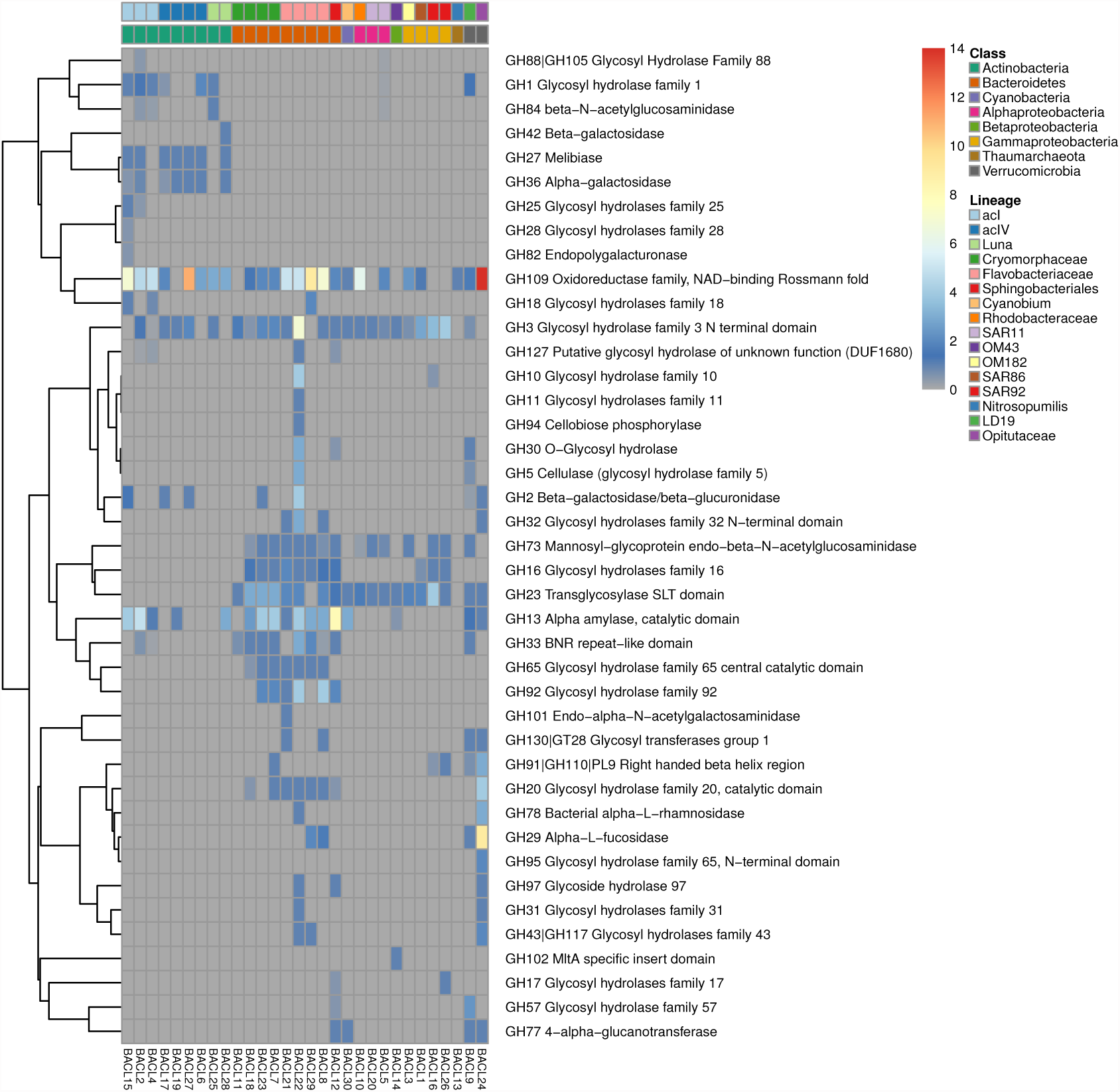
Abundance of glycoside hydrolase genes in MAG clusters. Counts were averaged over MAGs in each genome cluster.

**Supplementary Figure 7.**
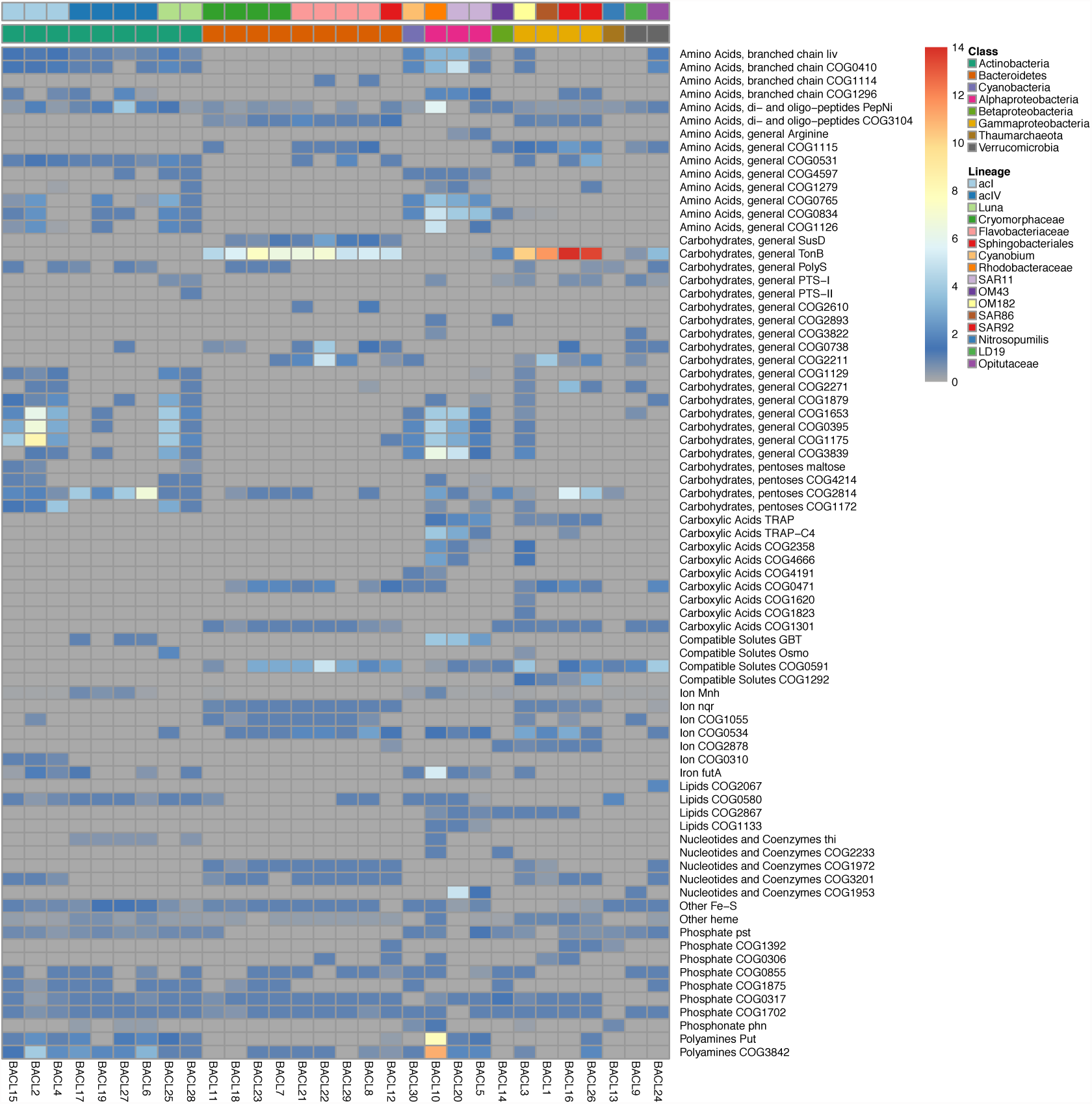
Abundance of transporters in MAG clusters. Counts of transporter genes were averaged over MAGs in each genome cluster and over transport types if applicable (see Supplementary Dataset 4).

**Supplementary Figure 8.**
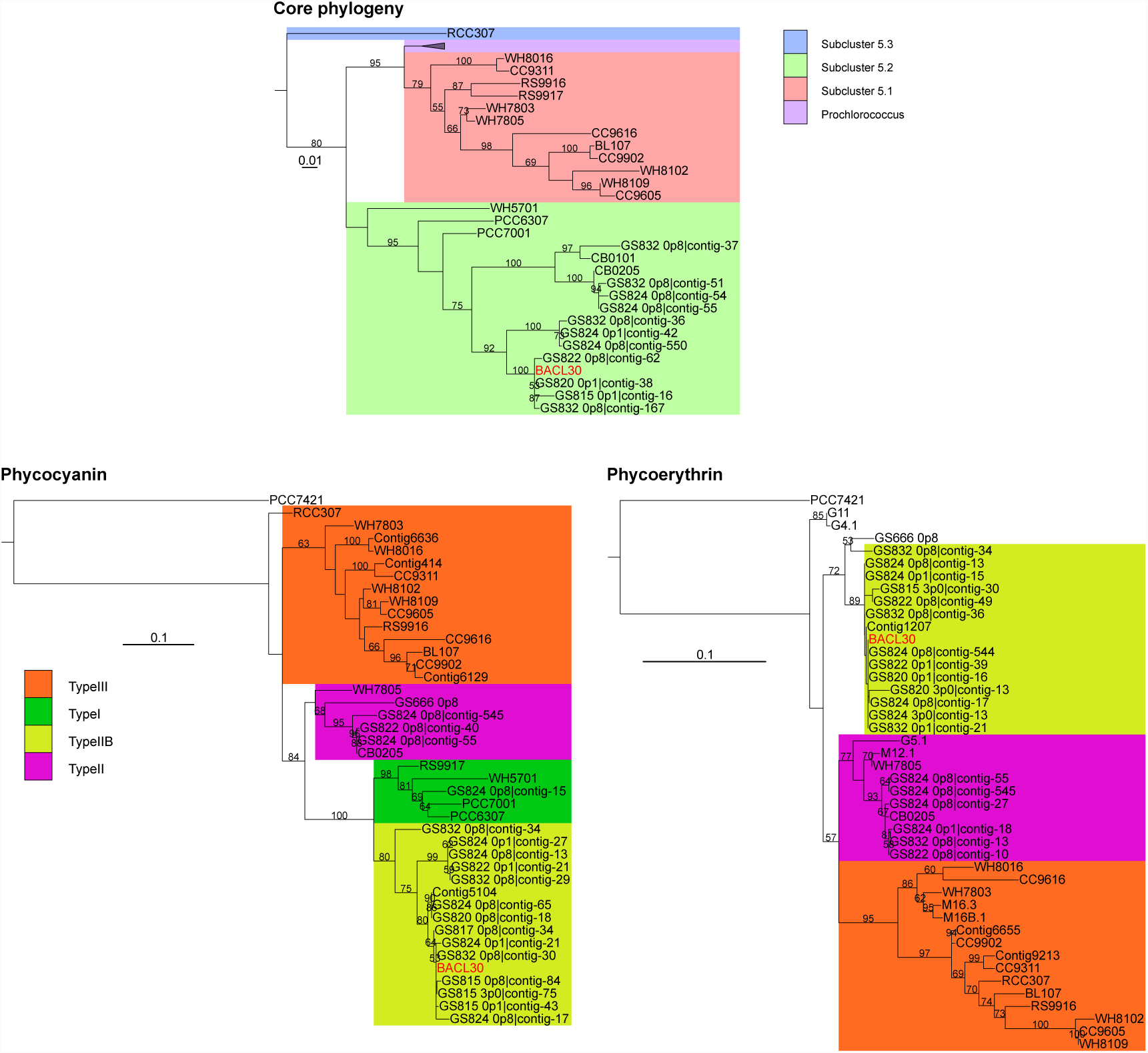
Core genome and pigment phylogeny of picocyanobacteria and BACL30. Phylogenetic trees are shown for core-genome proteins in picocyanobacteria (as in Larsson et al 2014) as well as concatenated alignments for phycocyanin (*cpcBA*) and phycoerythrin (*cpeBA*) gene products. In the core phylogeny, clade colors indicate subcluster designations for picocyanobacteria. In the pigment phylogenies, clade colors show pigment type designations. Scale bars indicate expected substitutions per site.

**Supplementary Figure 9.**
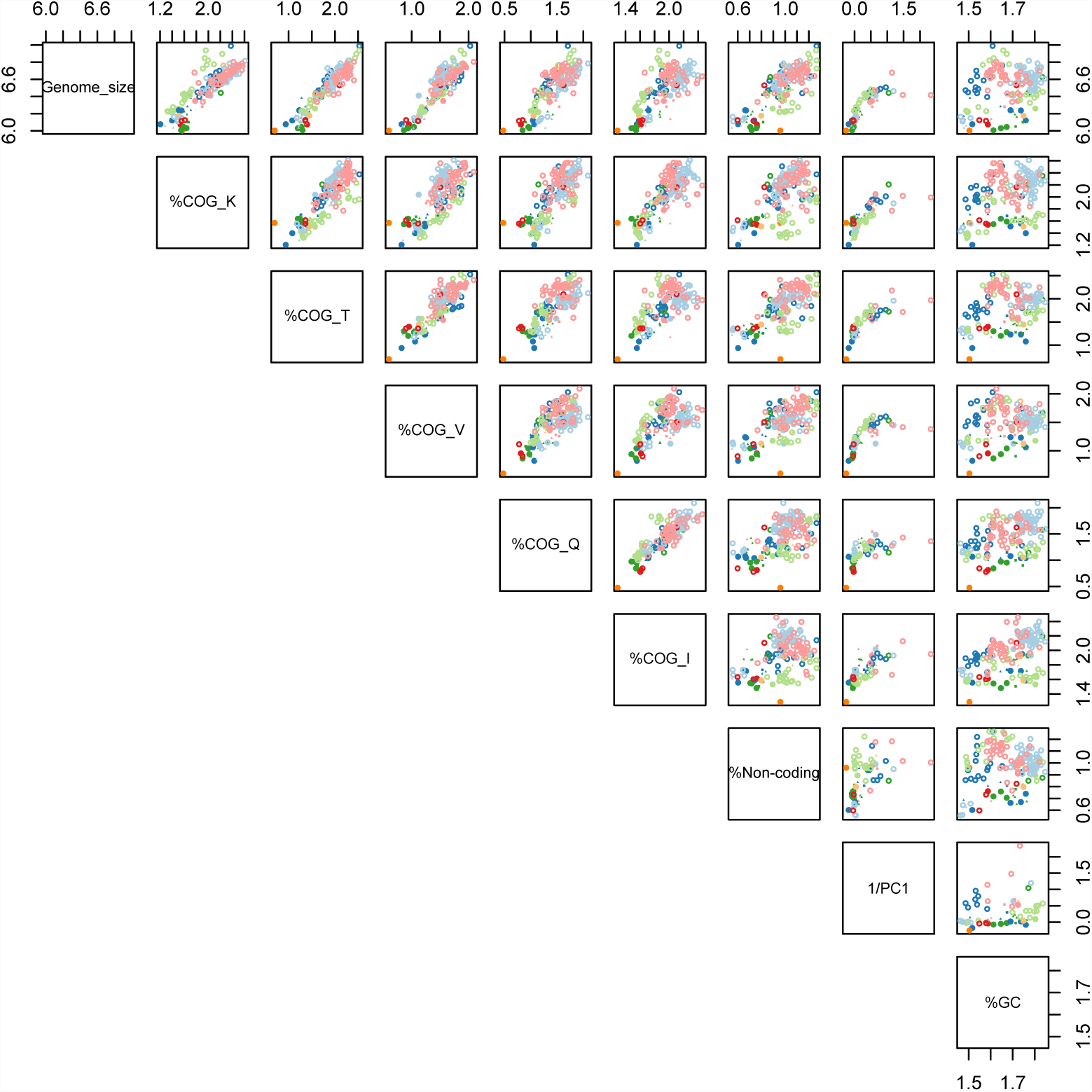
Pairwise scatter-plots showing correlations of genomes features among MAGs and isolate genomes. PC1 is the first principal component of the PCA of Figure 3A (that included all parameters except genome-size). Genomes are color-coded and shaped as in Figure 3.

**Supplementary Figure 10.**
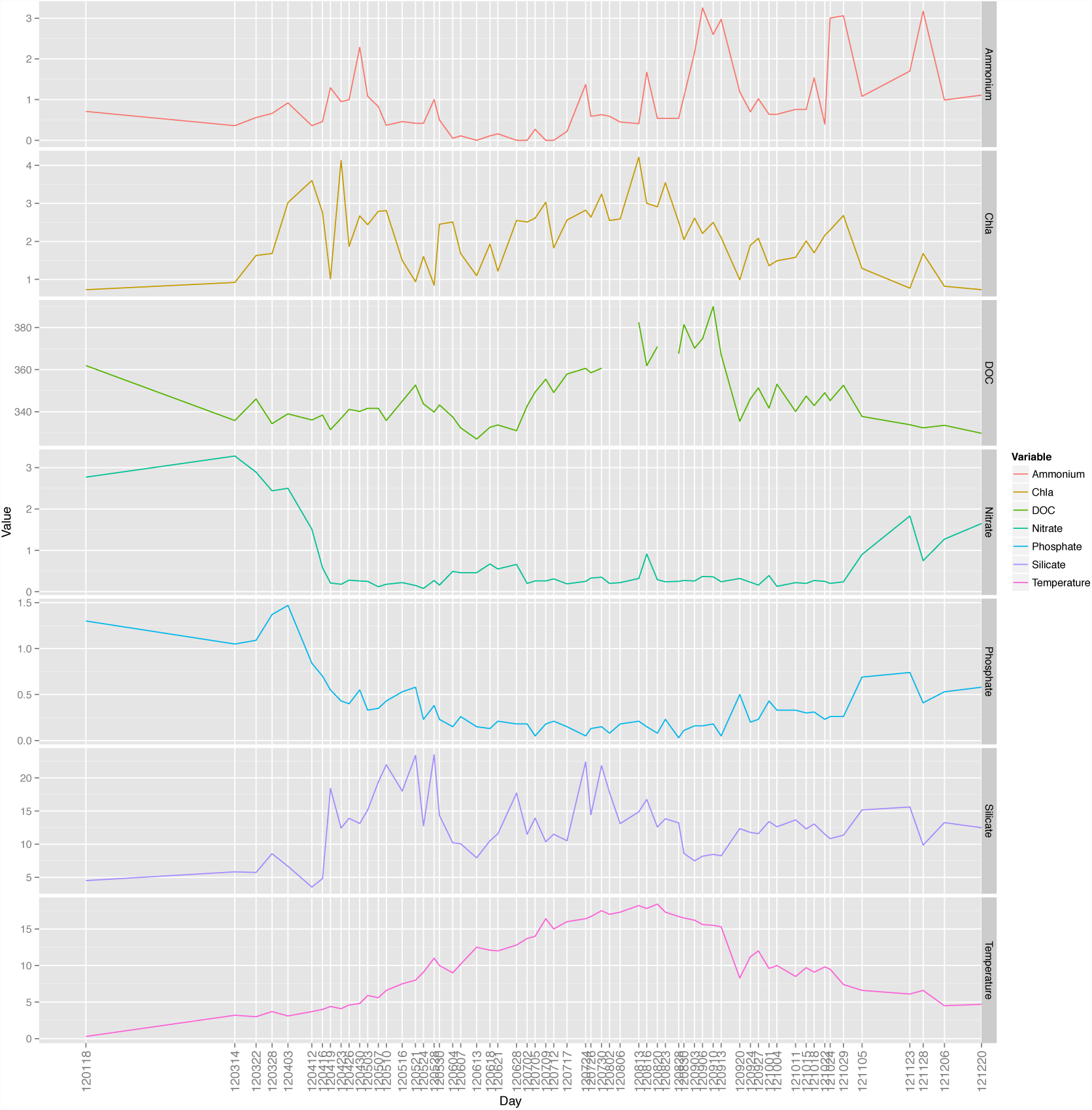
Variation in temperature (in ° C) and concentrations of nutrients, Chlorophyll a and DOC (in μg/L for Chlorophyll a and in μM for the rest).

**Supplementary Figure 11.**
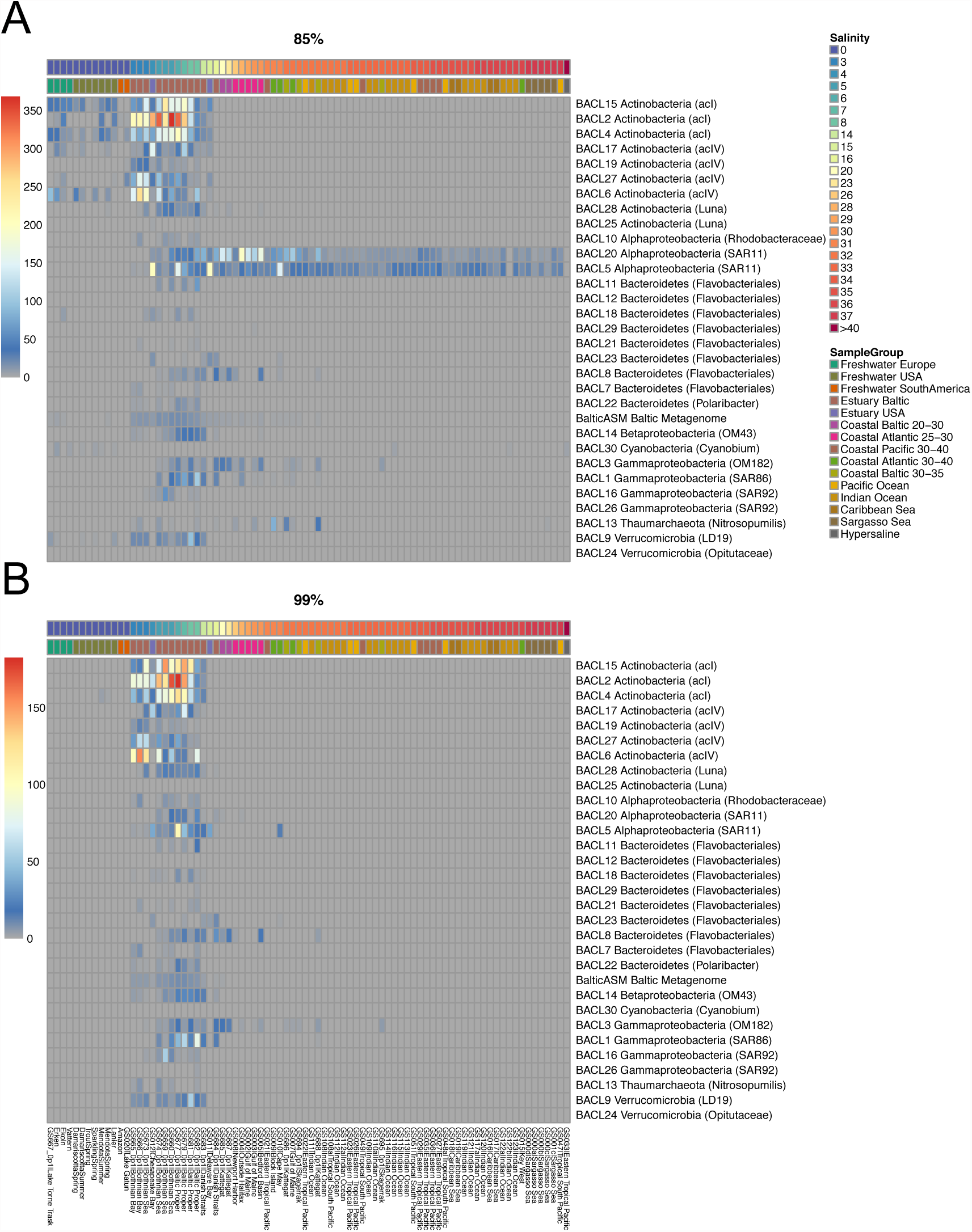
Biogeographical abundance profiles of MAGs. Heatmap plots showing the abundance of recruited reads from various samples and sample groups to each of the 30 MAG clusters at the 85% (A) and 99% (B) nucleotide identity cutoff levels. Shown values represent number of recruited reads/kb of genome per 10,000 queried reads. As in Figure 5 but with all samples in each sample group shown.

**Supplementary Figure 12.**
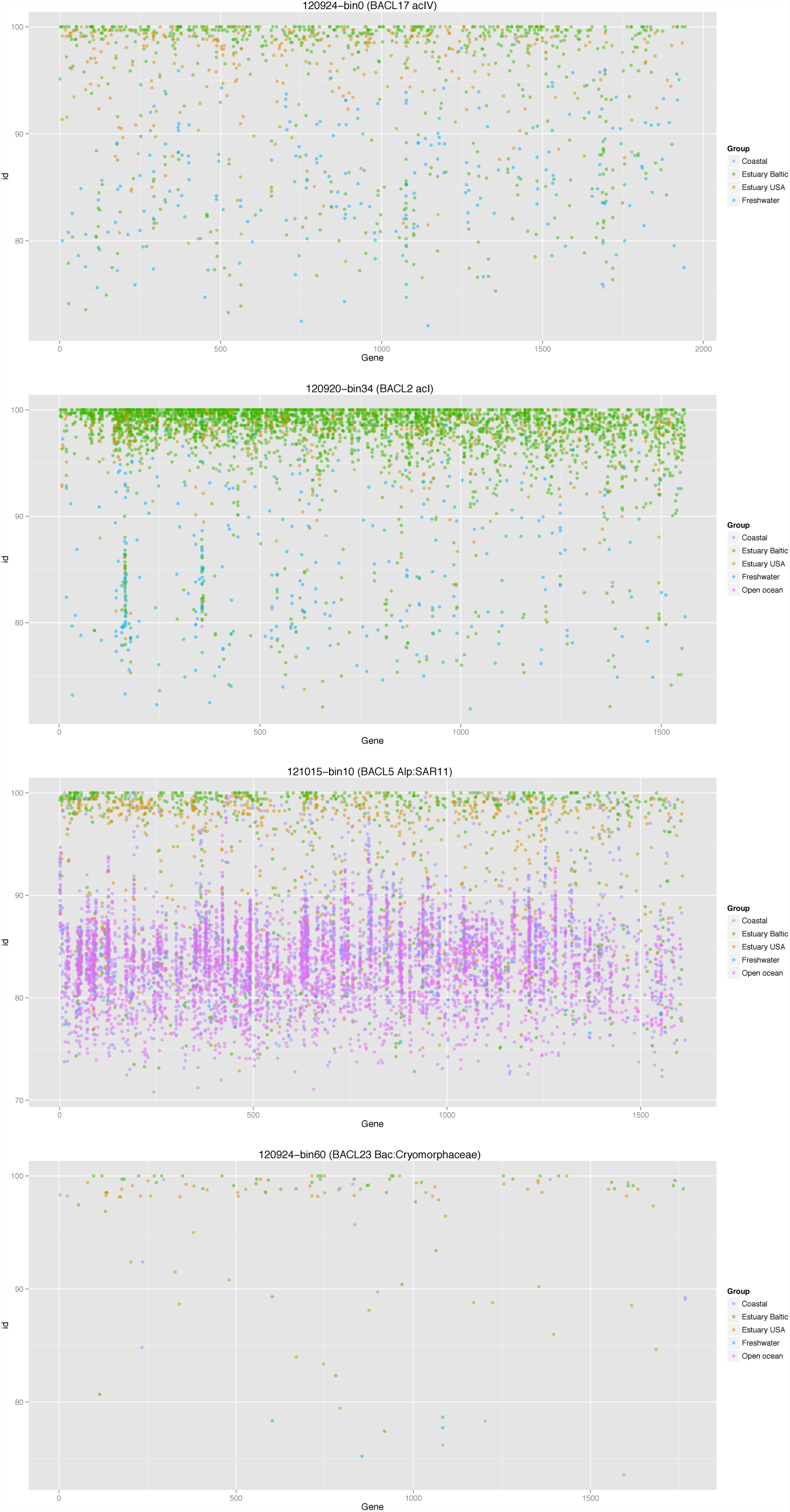
Fragment recruitment plots in selected MAG clusters. For each MAG cluster, the largest MAG is shown with the x-axis representing predicted open reading frames and the y-axis representing the nucleotide identity in %. Consequently, each recruited read is shown at a specific gene (x coordinate) and % identity (y coordinate) and further colored by sample group shown in the legend.

**Supplementary Table 1.** Results from ANOSIM analysis of clustering of MAGs based on functional annotations. For all analyses, phyla, classes or lineages with only 1 representative were removed.

| **Annotation** | **Phylum-p** | **Phylum-R** | **Class-p** | **Class-R** | **Lineage-p** | **Lineage-R** |
| --- | --- | --- | --- | --- | --- | --- |
| **TIGRFAM** | 0.001 | 0.874 | 0.001 | 0.966 | 0.001 | 0.973 |
| **PFAM** | 0.001 | 0.889 | 0.001 | 0.984 | 0.001 | 0.970 |
| **COG** | 0.001 | 0.868 | 0.001 | 0.967 | 0.001 | 0.977 |
| **EC** | 0.001 | 0.856 | 0.001 | 0.942 | 0.001 | 0.968 |
| **Minpath** | 0.001 | 0.785 | 0.001 | 0.840 | 0.001 | 0.984 |
| **Transporters** | 0.001 | 0.676 | 0.001 | 0.874 | 0.001 | 0.965 |

## References

1. Connon, S. A. & Giovannoni, S. J. High-Throughput Methods for Culturing Microorganisms in Very-Low-Nutrient Media Yield Diverse New Marine Isolates. Appl. Environ. Microbiol. 68, 3878–3885 (2002).

2. Lauro, F. M. et al. The genomic basis of trophic strategy in marine bacteria. Proc. Natl. Acad. Sci. U. S. A. 106, 15527–15533 (2009).

3. Swan, B. K. et al. Prevalent genome streamlining and latitudinal divergence of planktonic bacteria in the surface ocean. Proc. Natl. Acad. Sci. U. S. A. 110, 11463–11468 (2013).

4. Yooseph, S. et al. Genomic and functional adaptation in surface ocean planktonic prokaryotes. Nature 468, 60–66 (2010).

5. Tyson, G. W. et al. Community structure and metabolism through reconstruction of microbial genomes from the environment. Nature 428, 37–43 (2004).

6. Venter, J. C. et al. Environmental Genome Shotgun Sequencing of the Sargasso Sea. Science 304, 66–74 (2004).

7. Rusch, D. B. et al. The Sorcerer II Global Ocean Sampling Expedition: Northwest Atlantic through Eastern Tropical Pacific. PLoS Biol. DOI: 10.1371/journal.pbio.0050077 (2007).

8. Dupont, C. L. et al. Functional Tradeoffs Underpin Salinity-Driven Divergence in Microbial Community Composition. PLoS One DOI: 10.1371/journal.pone.0089549 (2014).

9. Martinez-Garcia, M. et al. High-throughput single-cell sequencing identifies photoheterotrophs and chemoautotrophs in freshwater bacterioplankton. ISMEJ 6, 113–123 (2011).

10. Ghylin, T. W. et al. Comparative single-cell genomics reveals potential ecological niches for the freshwater acI Actinobacteria lineage. ISMEJ 8, 2503–2516 (2014).

11. Wallner, G., Fuchs, B., Spring, S., Beisker, W. & Amann, R. Flow sorting of microorganisms for molecular analysis. Appl. Environ. Microbiol. 63, 4223–4231 (1997).

12. Woyke, T. et al. Assembling the marine metagenome, one cell at a time. PLoS One 4, e5299 (2009).

13. Yilmaz, S. & Singh, A. K. Single cell genome sequencing. Curr. Opin. Biotechnol. 23, 437–443 (2012).

14. Abe, T., Sugawara, H., Kinouchi, M., Kanaya, S. & Ikemura, T. Novel phylogenetic studies of genomic sequence fragments derived from uncultured microbe mixtures in environmental and clinical samples. DNA Res. 12, 281–290 (2005).

15. Dick, G. J. et al. Community-wide analysis of microbial genome sequence signatures. Genome Biol. 10, R85 (2009).

16. Sharon, I. et al. Time series community genomics analysis reveals rapid shifts in bacterial species, strains, and phage during infant gut colonization. Genome Res. 23, 111–120 (2012).

17. Albertsen, M. et al. Genome sequences of rare, uncultured bacteria obtained by differential coverage binning of multiple metagenomes. Nat. Biotechnol. 31, 533–538 (2013).

18. Imelfort, M. et al. GroopM: an automated tool for the recovery of population genomes from related metagenomes. PeerJ e603 (2014).

19. Nielsen, H. B. et al. Identification and assembly of genomes and genetic elements in complex metagenomic samples without using reference genomes. Nat. Biotechnol. 32, 822–828 (2014).

20. Alneberg, J. et al. Binning metagenomic contigs by coverage and composition. Nat. Methods 11, 1144–1146 (2014).

21. Ojaveer, H. et al. Status of Biodiversity in the Baltic Sea. PLoS One DOI: 10.1371/journal.pone.0012467 (2010).

22. Herlemann, D. P. R. et al. Transitions in bacterial communities along the 2000 km salinity gradient of the Baltic Sea. ISME J. 5, 1571–1579 (2011).

23. Andersson, A. F., Rieman, L. & Bertilsson, S. Pyrosequencing reveals contrasting seasonal dynamics of taxa within Baltic Sea bacterioplankton communities. ISMEJ 4, 171–181 (2010).

24. Lindh, M. V. et al. Disentangling seasonal bacterioplankton population dynamics by high frequency sampling. Environ. Microbiol. doi: 10.1111/1462-2920.12720 (2015).

25. Konstantinidis, K. T., Ramette, A. & Tiedje, J. M. The bacterial species definition in the genomic era. Philos. Trans. R. Soc. Lond. B Biol. Sci. 361, 1929–1940 (2006).

26. Herlemann, D. P. R. et al. Metagenomic De Novo Assembly of an Aquatic Representative of the Verrucomicrobial Class Spartobacteria. MBio 4, e00569–12 (2013).

27. Teeling, H. et al. Substrate-controlled succession of marine bacterioplankton populations induced by a phytoplankton bloom. Science 336, 608–611 (2012).

28. Tang, K., Jiao, N., Liu, K., Zhang, Y. & Li, S. Distribution and functions of TonB-dependent transporters in marine bacteria and environments: implications for dissolved organic matter utilization. PLoS One 7, e41204 (2012).

29. Alonso-Sáez, L. et al. Role for urea in nitrification by polar marine Archaea. Proc. Natl. Acad. Sci. U. S. A. 109, 17989–17994 (2012).

30. Giovannoni, S. J. et al. The small genome of an abundant coastal ocean methylotroph. Environ. Microbiol. 10, 1771–1782 (2008).

31. Walsh, D. A., Lafontaine, J. & Grossart, H.-P. in Lateral Gene Transfer in Evolution 55–77 (Springer New York, 2013).

32. Noinaj, N., Guillier, M., Barnard, T. J. & Buchanan, S. K. TonB-dependent transporters: regulation, structure, and function. Annu. Rev. Microbiol. 64, 43–60 (2010).

33. Iverson, V. et al. Untangling genomes from metagenomes: revealing an uncultured class of marine Euryarchaeota. Science 335, 587–590 (2012).

34. Di Rienzi, S. C. et al. The human gut and groundwater harbor non-photosynthetic bacteria belonging to a new candidate phylum sibling to Cyanobacteria. Elife 2, e01102 (2013).

35. Cho, J.-C. & Giovannoni, S. J. Cultivation and growth characteristics of a diverse group of oligotrophic marine Gammaproteobacteria. Appl. Environ. Microbiol. 70, 432–440 (2004).

36. Meon, B. & Kirchman, D. L. Dynamics and molecular composition of dissolved organic material during experimental phytoplankton blooms. Mar. Chem. 75, 185–199 (2001).

37. Oh, H.-M., Kang, I., Ferriera, S., Giovannoni, S. J. & Cho, J.-C. Genome sequence of the oligotrophic marine Gammaproteobacterium HTCC2143, isolated from the Oregon Coast. J. Bacteriol. 192, 4530–4531 (2010).

38. Zwart, G., Crump, B. C., Kamst-van Agterveld, M. P., Hagen, F. & Han, S. K. Typical freshwater bacteria: an analysis of available 16S rRNA gene sequences from plankton of lakes and rivers. Aquat. Microb. Ecol. 28, (2002).

39. Serkebaeva, Y. M., Kim, Y., Liesack, W. & Dedysh, S. N. Pyrosequencing-based assessment of the bacteria diversity in surface and subsurface peat layers of a northern wetland, with focus on poorly studied phyla and candidate divisions. PLoS One 8, e63994 (2013).

40. Dunfield, P. F. et al. Methane oxidation by an extremely acidophilic bacterium of the phylum Verrucomicrobia. Nature 450, 879–882 (2007).

41. Zhu, Y. et al. Staphylococcus aureus biofilm metabolism and the influence of arginine on polysaccharide intercellular adhesin synthesis, biofilm formation, and pathogenesis. Infect. Immun. 75, 4219–4226 (2007).

42. Welander, P. V. et al. Hopanoids play a role in membrane integrity and pH homeostasis in Rhodopseudomonas palustris TIE-1. J. Bacteriol. 191, 6145–6156 (2009).

43. Newton, R. J., Jones, S. E., Eiler, A., McMahon, K. D. & Bertilsson, S. A guide to the natural history of freshwater lake bacteria. Microbiol. Mol. Biol. Rev. 75, 14–49 (2011).

44. Larsson, J. et al. Picocyanobacteria containing a novel pigment gene cluster dominate the brackish water Baltic Sea. ISMEJ 8, 1892–1903 (2014).

45. Palenik, B. et al. Genome sequence of Synechococcus CC9311: Insights into adaptation to a coastal environment. Proc. Natl. Acad. Sci. U. S. A. 103, 13555–13559 (2006).

46. Giovannoni, S. J., Thrash, J. C. & Temperton, B. Implications of streamlining theory for microbial ecology. ISMEJ 8, 1553–1565 (2014).

47. Dufresne, A. et al. Genome sequence of the cyanobacterium Prochlorococcus marinus SS120, a nearly minimal oxyphototrophic genome. Proc. Natl. Acad. Sci. U. S. A. 100, 10020–10025 (2003).

48. Giovannoni, S. J. et al. Genome streamlining in a cosmopolitan oceanic bacterium. Science 309, 1242–1245 (2005).

49. Button, D. K. Nutrient uptake by microorganisms according to kinetic parameters from theory as related to cytoarchitecture. Microbiol. Mol. Biol. Rev. 62, 636–645 (1998).

50. Nedashkovskaya, O. I. et al. Polaribacter butkevichii sp. nov., a novel marine mesophilic bacterium of the family Flavobacteriaceae. Curr. Microbiol. 51, 408–412 (2005).

51. Gómez-Pereira, P. R. et al. Genomic content of uncultured Bacteroidetes from contrasting oceanic provinces in the North Atlantic Ocean. Environ. Microbiol. 14, 52–66 (2012).

52. Wong, E. et al. The Vibrio cholerae colonization factor GbpA possesses a modular structure that governs binding to different host surfaces. PLoS Pathog. 8, e1002373 (2012).

53. Sabath, N., Ferrada, E., Barve, A. & Wagner, A. Growth temperature and genome size in bacteria are negatively correlated, suggesting genomic streamlining during thermal adaptation. Genome Biol. Evol. 5, 966–977 (2013).

54. Parveen, B. et al. Diversity and dynamics of free-living and particle-associated Betaproteobacteria and Actinobacteria in relation to phytoplankton and zooplankton communities. FEMS Microbiol. Ecol. 77, 461–476 (2011).

55. Penn, K., Wang, J., Fernando, S. C. & Thompson, J. R. Secondary metabolite gene expression and interplay of bacterial functions in a tropical freshwater cyanobacterial bloom. ISME J. 8, 1866–1878 (2014).

56. Kuo, C. & Ochman, H. Inferring clocks when lacking rocks: the variable rates of molecular evolution in bacteria. Biol. Direct 4, doi:10.1186/1745-6150 (2009).

57. Coolen, M. & Overmann, J. 217 000-year-old DNA sequences of green sulfur bacteria in Mediterranean sapropels and their implications for the reconstruction of the paleoenvironment. Environ. Microbiol. 9, 238–249 (2007).

58. Logares, R. et al. Infrequent marine–freshwater transitions in the microbial world. Trends Microbiol. 17, 414–422 (2009).

59. Nilsson, J. et al. Matrilinear phylogeography of Atlantic salmon (Salmo salar L.) in Europe and postglacial colonization of the Baltic Sea area. Mol. Ecol. 10, 89–102 (2001).

60. Luttikhuizen, P. C., Drent, J. & Baker, A. J. Disjunct distribution of highly diverged mitochondrial lineage clade and population subdivision in a marine bivalve with pelagic larval dispersal. Mol. Ecol. 12, 2215–2229 (2003).

61. Johannesson, K. & André, C. Life on the margin: genetic isolation and diversity loss in a peripheral marine ecosystem, the Baltic Sea. Mol. Ecol. 15, 2013–2029 (2006).

62. Martin, M. Cutadapt removes adapter sequences from high-throughput sequencing reads. Bioinformatics in Action 17, 10–12 (2012).

63. Xu, H. et al. FastUniq: A Fast De Novo Duplicates Removal Tool for Paired Short Reads. PLoS One 7, e52249 (2012).

64. Boisvert, S., Laviolette, F. & Corbeil, J. Ray: Simultaneous Assembly of Reads from a Mix of High-Throughput Sequencing Technologies. J. Comput. Biol. 17, 1519–1533 (2010).

65. Langmead, B. & Salzberg, S. L. Fast gapped-read alignment with Bowtie 2. Nat. Methods 9, 357–359 (2012).

66. Li, H. et al. The Sequence Alignment/Map format and SAMtools. Bioinformatics 25, 1078–1079 (2009).

67. Quinlan, A. R. & Hall, I. M. BEDTools: a flexible suite of utilities for comparing genomic features. Bioinformatics 26, 841–842 (2010).

68. Hyatt, D., Locascio, P. F., Hauser, L. J. & Uberbacher, E. C. Gene and translation initiation site prediction in metagenomic sequences. Bioinformatics 28, 2223–2230 (2012).

69. Kurtz, S. et al. Versatile and open software for comparing large genomes. Genome Biol. 5, R12 (2004).

70. Seemann, T. Prokka: rapid prokaryotic genome annotation. Bioinformatics 30, 2068–2069 (2014).

71. Ye, Y. & Doak, T. G. A Parsimony Approach to Biological Pathway Reconstruction/Inference for Genomes and Metagenomes. PLoS Comput. Biol. 5, e1000465 (2009).

72. Caspi, R. et al. The MetaCyc database of metabolic pathways and enzymes and the BioCyc collection of pathway/genome databases. Nucleci Acids Research 40, D742–53 (2012).

73. Parks, D. H., Tyson, G. W., Hugenholtz, P. & Beiko, R. G. STAMP: statistical analysis of taxonomic and functional profiles. Bioinformatics 30, 3123–3124 (2014).

74. Darling, A. E. et al. PhyloSift: phylogenetic analysis of genomes and metagenomes. PeerJ 2, e243 (2014).

75. Wu, S., Zhu, Z., Fu, L., Niu, B. & Li, W. WebMGA: a customizable web server for fast metagenomic sequence analysis. BMC Genomics 12, 444 (2011).

76. Kopylova, E., Noé, L. & Touzet, H. SortMeRNA: fast and accurate filtering of ribosomal RNAs in metatranscriptomic data. Bioinformatics 28, 3211–3217 (2012).

77. Pruesse, E., Peplies, J. & Glöckner, F. O. SINA: accurate high troughput multiple sequence alignment of ribosomal RNA genes. Bioinformatics doi: 10.1093/bioinformatics/bts252(2012).

78. Segata, N., Börnigen, D., Morgan, X. C. & Huttenhower, C. PhyloPhlAn is a new method for improved phylogenetic and taxonomic placement of microbes. Nat. Commun. 4, 2304 (2013).

79. Letunic, I. & Bork, P. Interactive Tree Of Life v2: online annotation and display of phylogenetic trees made easy. Nucleic Acids Res. 39, W475–8 (2011).

80. Auch, A. F., Klenk, H.-P. & Göker, M. Standard operating procedure for calculating genome-to-genome distances based on high-scoring segment pairs. Stand. Genomic Sci. 2, 142–148 (2010).

81. Eiler, A. et al. Unveiling distribution patterns of freshwater phytoplankton by a next generation sequencing based approach. PLoS One 8, e53516 (2013).

82. Oh, S. et al. Metagenomic Insights into the Evolution, Function, and Complexity of the Planktonic Microbial Community of Lake Lanier, a Temperate Freshwater Ecosystem. Appl. Environ. Microbiol. 77, 6000–6011 (2011).

83. Ghai, R. et al. Metagenomics of the Water Column in the Pristine Upper Course of the Amazon River. PLoS One 6, e23785 (2011).

84. Kang, I. et al. Genome sequence of “Candidatus Aquiluna” sp. strain IMCC13023, a marine member of the Actinobacteria isolated from an arctic fjord. J. Bacteriol. 194, 3550–3551 (2012).

85. Hahn, M. W., Schmidt, J., Taipale, S. J., Doolittle, W. F. & Koll, U. Rhodoluna lacicola gen. nov., sp. nov., a planktonic freshwater bacterium with stream-lined genome. Int. J. Syst. Evol. Microbiol. 64, 3254–3263 (2014).

86. Garcia, S. L. et al. Metabolic potential of a single cell belonging to one of the most abundant lineages in freshwater bacterioplankton. ISME J. 7, 137–147 (2013).

87. Glöckner, F. O. et al. Comparative 16S rRNA Analysis of Lake Bacterioplankton Reveals Globally Distributed Phylogenetic Clusters Including an Abundant Group of Actinobacteria. Appl. Environ. Microbiol. 66, 5053–5065 (2000).

88. Newton, R. J., Kent, A. D., Triplett, E. W. & McMahon, K. D. Microbial community dynamics in a humic lake: differential persistence of common freshwater phylotypes. Environ. Microbiol. 8, 956–970 (2006).

89. Wu, X., Xi, W., Ye, W. & Yang, H. Bacterial community composition of a shallow hypertrophic freshwater lake in China, revealed by 16S rRNA gene sequences. FEMS Microbiol. Ecol. 61, 85–96 (2007).

90. Humbert, J.-F. et al. Comparison of the structure and composition of bacterial communities from temperate and tropical freshwater ecosystems. Environ. Microbiol. 11, 2339–2350 (2009).

91. Bowman, J. in The Prokaryotes (eds. Rosenberg, E., DeLong, E., Lory, S., Stackebrandt,E. & Thompson, F.) 539–550 (Springer Berlin Heidelberg, 2014).

92. Huggett, M. J., Hayakawa, D. H. & Rappé, M. S. Genome sequence of strain HIMB624, a cultured representative from the OM43 clade of marine Betaproteobacteria. Stand. Genomic Sci. 6, 11–20 (2012).

93. Crosbie, N. D., Pöckl, M. & Weisse, T. Dispersal and phylogenetic diversity of nonmarine picocyanobacteria, inferred from 16S rRNA gene and cpcBA-intergenic spacer sequence analyses. Appl. Environ. Microbiol. 69, 5716–5721 (2003).

94. Hugenholtz, P., Pitulle, C., Hershberger, K. L. & Pace, N. R. Novel division level bacterial diversity in a Yellowstone hot spring. J. Bacteriol. 180, 366–376 (1998).

95. Zwart, G. et al. Divergent members of the bacterial division Verrucomicrobiales in a temperate freshwater lake. FEMS Microbiol. Ecol. 25, 159–169 (1998).

96. Op den Camp, H. J. et al. Environmental, genomic and taxonomic perspectives on methanotrophic Verrucomicrobia. Environ. Microbiol. Rep. 1, (2009).

97. Yoon, J. et al. Coraliomargarita akajimensis gen. nov., sp. nov., a novel member of the phylum “Verrucomicrobia” isolated from seawater in Japan. IJSEM 57, 959–963 (2007).

98. Karner, M. B., DeLong, E. F. & Karl, D. M. Archaeal dominance in the mesopelagic zone of the Pacific Ocean. Nature 409, 507–510 (2001).

99. Wuchter, C. et al. Archaeal nitrification in the ocean. Proc. Natl. Acad. Sci. U. S. A. 103, 12317–12322 (2006).

100. Walker, C. B. et al. Nitrosopumilus maritimus genome reveals unique mechanisms for nitrification and autotrophy in globally distributed marine crenarchaea. Proc. Natl. Acad. Sci. U. S. A. 107, 8818–8823 (2010).

101. Park, S. et al. Genomes of Two New Ammonia-Oxidizing Archaea Enriched from Deep Marine Sediments. PLoS One DOI: 10.1371/journal.pone.0096449 (2014).

102. Labrenz, M. et al. Relevance of a crenarchaeotal subcluster related to Candidatus Nitrosopumilus maritimus to ammonia oxidation in the suboxic zone of the central Baltic Sea. ISMEJ 4, 1496–1508 (2010).

103. Pruitt, K. D. et al. RefSeq: an update on mammalian reference sequences. Nucleic Acids Res. 42, D756–63 (2014).

